# Duration of Initial Viremia Modulates Functional Properties of HIV-specific T Cell Receptors

**DOI:** 10.64898/2026.01.29.702605

**Authors:** Funsho J. Ogunshola, Nishant K. Singh, Vincent Butty, Anurag R. Mishra, Zacharia Habte, Liza Vecchiarello, Ahmed Fahad, Kate B. Juergens, Sophia Cheever, Marius Allombert, Anabelle Webber, Alicja Piechocka-Trocha, Nasreen Ismail, Anusha Nathan, Xiaolong Li, Kavidha Reddy, Kamini Gounder, Omolara O. Baiyegunhi, David R. Collins, Musie Ghebremichael, Gaurav Gaiha, Krista Dong, Brandon J. Dekosky, Thumbi Ndung’u, Michael E. Birnbaum, Bruce D. Walker

## Abstract

Virus-specific CD8^+^ T cells are crucial in controlling chronic human viral infections such as HIV-1, but the effect of persistent antigen exposure on T cell repertoire formation is not well understood. In this study, we examined epitope-specific CD8^+^ T cell repertoires in people living with HIV-1, where duration of viremia following hyperacute infection was modulated by the time of initiation of continuous suppressive antiretroviral therapy (ART). After ART-induced undetectable viremia in persons expressing the same HLA class I allele, we analyzed the impact of early (n=6) versus delayed (n=6) ART initiation on the clonotypic composition, clonotypic cross-reactivity, functional avidity and memory differentiation profile of the HIV-specific T cell repertoire restricted by HLA-B*58:01. Using a panel of barcoded tetramers, we mapped T cell receptor (TCR) clonotypes specific for three dominant epitopes and their variants. Both groups exhibited polyclonal TCR repertoires with evidence of cross-reactivity, which was significantly enriched in donors with prolonged antigen exposure. Within this cohort, broadly cross-reactive clonotypes capable of recognizing all autologous variants were identified, but these were rare (<1%). Early ART initiation preserved repertoires characterized by higher-avidity TCRs and a relative enrichment of transitional memory CD8^+^ T cell subsets. These functional differences were not associated with differences in TRBV gene sharing, indicating that ART timing shapes repertoire quality and memory differentiation without altering TRBV gene bias. These findings demonstrate how antigen suppression dynamics differentially shape the breadth, functional sensitivity, and memory composition of the HIV-specific TCR repertoire, with implications for T cell-directed immunotherapies and HIV cure strategies.

**One Sentence Summary:** The duration of viral antigen exposure during early HIV infection shapes the functional quality, breadth, and memory composition of virus-specific CD8⁺ T cell receptor repertoires.

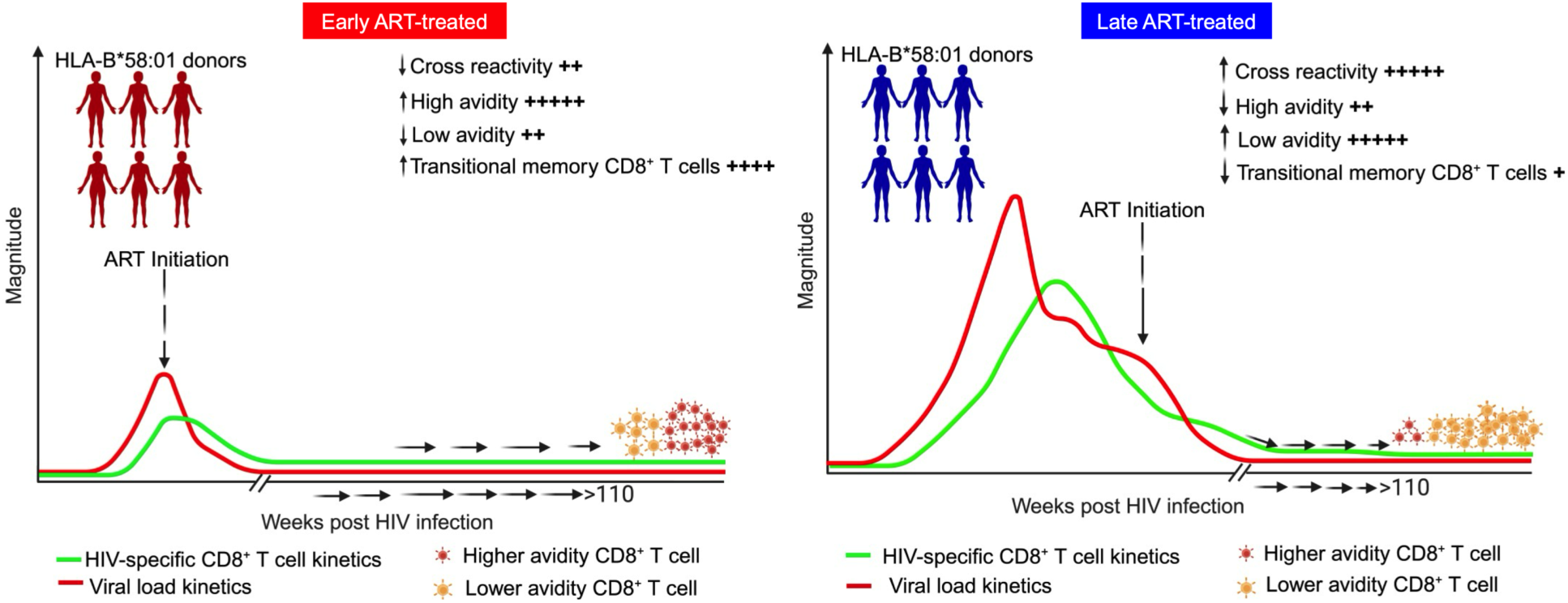

## Introduction

More than four decades since the identification of human immunodeficiency virus type 1 (HIV-1) as the cause of AIDS, there remains no effective vaccine^1–3^. Vaccines that focus on eliciting antibody responses have fallen short of protecting against infection, primarily due to rapid viral evolution and the need for extensive somatic hypermutation to achieve antibody effectiveness^4^. In contrast, cytotoxic CD8^+^ T lymphocytes (CTLs) offer a promising alternative for effectively controlling rapidly evolving viruses once infection occurs^5,6^.

In individuals who durably control HIV-1 without medications, which has been termed a functional cure^7^, HIV-specific CTLs have been shown to preferentially target conserved, mutationally constrained epitopes presented by Human Leukocyte Antigen (HLA) class I molecules, thereby reducing the likelihood of viral escape^8^. The importance of these CTLs in viral control is underscored by the strong association between certain HLA class I alleles and immune-mediated control of chronic viral infections^8,9^. Additionally, studies in rhesus macaques have demonstrated that depletion of CD8^+^ T cells results in an inability to control simian immunodeficiency virus (SIV) replication^10,11^, highlighting the critical role of CD8^+^ T cells as an antiviral host defense.

Several factors have been reported to influence the functional qualities of antigen-specific CTLs. The duration of antigen presentation by HLA molecules plays a pivotal role in determining the magnitude of T-cell responses during priming^12,13^. Limiting antigenic stimulation by approaches including pharmacologic elimination of genetically susceptible dendritic cells^12^, targeting peptide-HLA (pHLA) presentation with monoclonal antibodies^14^, or timed elimination of an infectious source of antigen by the administration of antibiotics^15^ reduces the magnitude of induced responses. However, the precise influence of this duration on the antigen-specific T cell receptor (TCR) clonotypic repertoire composition and function remains poorly understood.

A key factor in determining the quality of antigen-specific T cells is antigen sensitivity or functional avidity, which predicts the cell’s ability to respond to specific concentrations of its target antigen^16–20^. T cells with higher antigen sensitivity typically exhibit superior functionality^21^. The affinity or the strength of the interaction between the TCR and the pHLA complex modulates antigen sensitivity^13^. High-affinity TCRs are more likely to receive strong and sustained activation signals, often leading to their dominance in antigen-specific CD8^+^ T cell responses^18,22^. Moreover, the dynamics of TCR binding to the pHLA complex can influence processes such as clonal apoptosis and immune senescence^23,24^, impacting both the magnitude and quality of the T cell response. For example, in cytomegalovirus (CMV) infection, prolonged antigen exposure can cause a shift in TCR hierarchies, with low-affinity clones eventually overtaking high-affinity ones^24^. However, it remains unclear to what extent differences in the initial duration of antigen exposure impact TCR repertoire properties of antigen-reactive CD8^+^ T cells.

Here, we investigated how viremia in a chronic human viral infection influences TCR repertoire composition, cross-reactivity, avidity, and memory subsets in a unique cohort of persons identified with hyperacute HIV-1 infection^25–28^. In these persons with HIV (PWH), antigen exposure was modulated by immediate or delayed antiretroviral therapy (ART) initiation after HIV-1 detection, based on treatment guidelines at enrollment^29^. Cryopreserved peripheral blood mononuclear cell (PBMC) samples from individuals provided the opportunity to examine the impact of duration of antigen exposure on the clonotypic characteristics of the induced HIV-specific CD8^+^ T cells.

Study participants were recruited from the FRESH (Females Rising through Education, Support and Health) cohort, which focuses on identifying hyperacute HIV-1 infection before the initial peak in viremia^25^. This cohort, from a region of South Africa where the incidence of HIV infection was approximately 8-10% per year when the cohort was initiated, was designed with two main objectives: first, to provide women ages 18-23 years old and at high risk of HIV-1 infection with a pathway out of socioeconomic constraints through an intensive curriculum that included training in life skills and job readiness, as well as intensive HIV-1 prevention education, pre-exposure prophylaxis and ultimately employment opportunities; and second, to detect hyperacute HIV-1 infection by screening participants twice weekly using finger-prick blood samples for HIV-1 RNA. Importantly, to control for confounding HLA diversity, we focused on twelve persons all expressing the relatively protective *HLA-B*58:01* genotype^30,31^. Differences in the timing of ART initiation allowed for stratification by duration of detectable viremia

Our findings revealed limited broadly cross-reactive TCRs in both early and late treated groups, with HIV-specific CD8^+^ T cell clonotypes exhibiting lower avidity and the relative depletion of transitional memory populations following prolonged antigen exposure before ART initiation. These avidity profiles were independent of specific V gene usage. Molecular analysis further supported these observations, indicating that prolonged antigen exposure is associated with the generation of a lower-avidity HIV-specific CD8^+^ T cell repertoire compared to individuals treated during hyperacute HIV-1 infection.

## Results

### Modulation of HIV antigenemia by ART initiation in persons expressing HLA-B*58:01

Participants were selected based on diagnosis of acute infection and the expression of HLA-B*58:01 (**Supplementary Table 1**). This allele was selected because it was sufficiently represented in southern Africa^32,33^ and in our cohort to allow us match all participants at a single HLA class I allele. Viral load and CD4^+^ T cell counts were monitored longitudinally before and after ART initiation (**Figure 1**). The total duration of antigen exposure was defined as the number of days from initial detection of HIV-1 infection to the achievement of an undetectable viral load following ART initiation.

**Figure 1:**
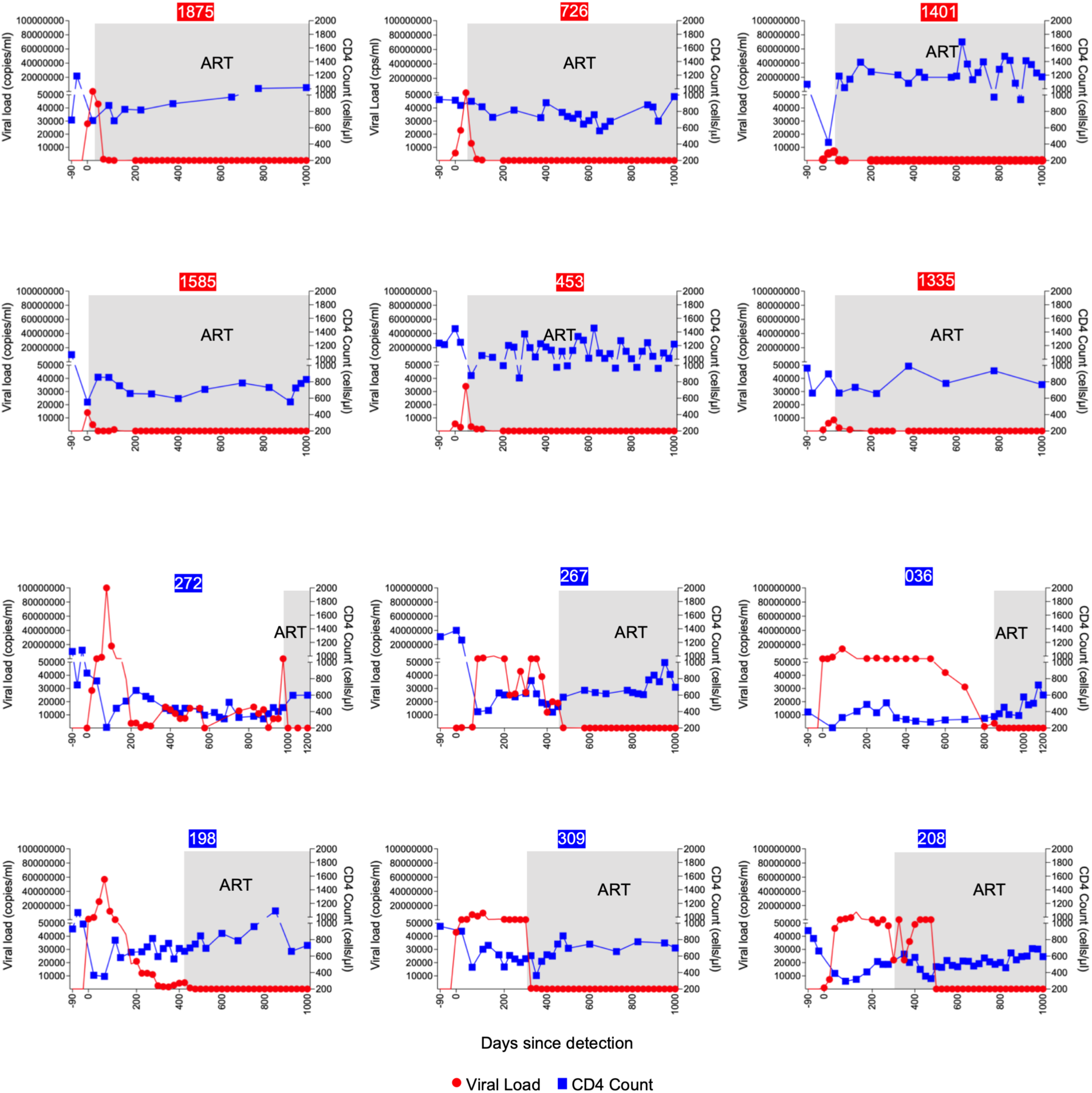
Longitudinal dynamics of CD4^+^ T cell counts and viral loads in HLA-B*58:01 participants. Longitudinal analysis of plasma HIV-1 RNA (red line) and absolute CD4^+^ T cell counts (blue line) in 12 participants identified in hyperacute infection and matched for the HLA-B*58:01 allele. The top two rows represent early treated individuals, while the bottom two rows correspond to those who initiated ART later. Gray shading indicates the periods during which participants were on intensive ART. Numbers in red boxes indicate early-treated participants; those in blue boxes are late-treated participants.

Each participant was identified at Fiebig stage I or II of HIV-1 infection^34^, before the time of peak viremia (**Table 1**). Those in the limited antigen exposure group (n=6) initiated ART immediately, following adoption of universal test-and-treat guidelines^35^. Peak viral loads ranged from 6,800-86,000 RNA copies/ml plasma, and undetectable viral loads were achieved after a median of 21 days following initiation of ART (range: 7 to 38 days). Participants in the prolonged antigen exposure group (n=6) were recruited before implementation of the immediate treatment policy, beginning ART after the CD4^+^ T cell count dropped below 350 cells/µl blood, in accordance with national treatment guidelines at the time. In this group, the median peak viral load was more than 1000-fold higher, ranging from 4 million to greater than 100 million RNA copies/ml, and the median duration of detectable plasma viremia was 684 days (range: 395 to 993 days), 30-fold longer. (**Table 2**).

**Table 1.**
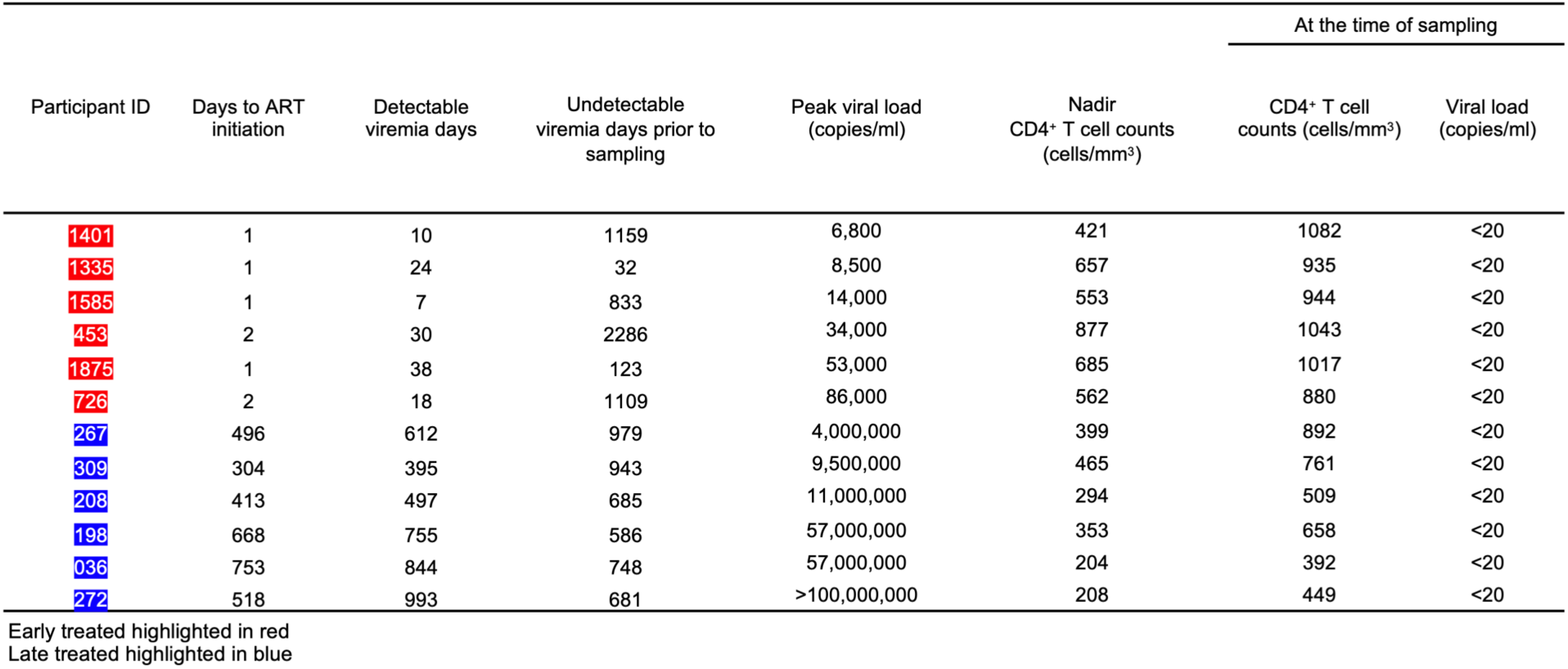
Demographic and clinical characteristics of the study participants.

**Table 2.** Summary of demographic and clinical characteristics of the study participants.

| Study cohort | Median Age (yr) | Ethnicity | Median detectable viremia days | Median undetectable viremia days prior to sampling | Median Nadir CD4 counts, (cells/mm<sup>3</sup>) | Median Peak viral load (RNA copies/ml) |
| --- | --- | --- | --- | --- | --- | --- |
| Early treated Participants (n = 6) | 21 (19-22) | Black Africans (100%) | 21 (7-38) | 971 (32-2286) | 609 (421-877) | 24000 (6800-86000) |
| Late treated Participants (n = 6) | 21 (18-22) | Black Africans (100%) | 683 (395-993) | 716 (586-979) | 323 (204-465) | 34X10<sup>6</sup> (4-100X10<sup>6</sup>) |

All participants were female

All participants selected targeted at least one of five previously described HLA-B*58:01-restricted HIV clade C epitopes (TW10_Gag(240-249)_, IW9_Pol(530-538)_, KW11_Env(59-69)_, RY11_Gag(76-86)_, and HW9_Nef(116-124)_) by *ex vivo* interferon-gamma (IFN-γ) Enzyme-Linked Immunospot (ELISPOT) assay during the acute phase of HIV infection (**Figure S1**). Due to sample availability, ELISPOT assay timing varied among participants, ranging from the second week following HIV-1 detection up to 24 weeks post detection of infection. All the participants were tested during the period of detectable viremia, except for one individual who was screened immediately after ART-mediated viral suppression (**see Table 1**).

Recognition of the Gag-derived TW10 epitope was the most frequent, detected *ex vivo* in nine of twelve participants. Magnitudes of responses were quite variable, and there was no consistently immunodominant epitope. The weakest overall responses were to to RY11_Gag(76-86)_ and HW9_Nef(116-124)_; due to limited sample availability, these were omitted from further analysis. Overall, these results revealed considerable heterogeneity in both magnitude and breadth of detectable *ex vivo* responses restricted by HLA-B*58:01 in the acute phase of infection, with only two of the twelve participants (1875 and 726) targeting all five epitopes, albeit at very different magnitudes. There were no significant differences in the breadth or magnitude of responses in the two groups at the time of sampling.

### Most consensus and variant HLA-B*58:01-restricted epitope peptides identified in acute infection effectively stabilize HLA class I

To determine T cell repertoire targeting not just consensus but all autologous HLA-B*58:01-restricted epitopes, we first sequenced the infecting virus in each of the twelve participants to determine autologous epitope variants. We focused on KW11, TW10 and IW9, which had been most dominantly targeted in the ELISPOT assays. Viral sequencing revealed heterogeneity in the dominant autologous epitope sequences, with several variants recurring in more than one participant (**Table 3 and Supplementary Table of Extended Epitope Sequences)**. Many of these variants have been previously documented in the Los Alamos HIV Immunology Database^36^.

**Table 3.**
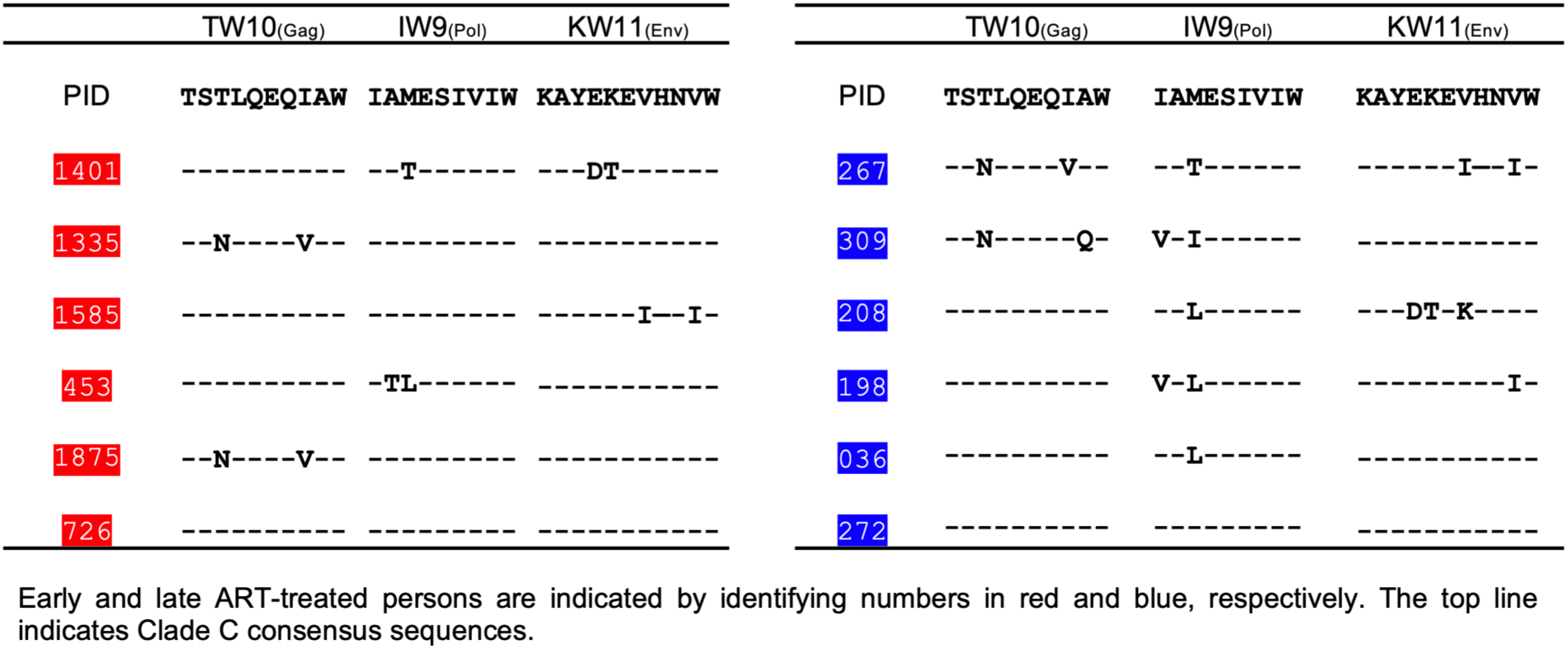
HLA-B*58:01 epitope sequences identified in participants during acute phase of HIV-1 infection.

We next determined the impact of viral variation on epitope presentation by evaluating the ability of HLA-B*58:01 to present these three epitopes and their variants using an *in vitro* peptide-HLA (pHLA) stability assay using TAP-deficient, mono-allelic HLA-B*58:01 class I-expressing cell lines^37^. This assay measures the stabilization of HLA class I surface expression upon loading with exogenous peptides. As a positive control, we included a well-characterized HLA-B*58:01-restricted Epstein-Barr virus (EBV) VW9_LMP1(125-133)_ peptide^38^, as well as the HLA-B*81:01-restricted HIV TL9_Gag(180-188)_ peptide^39^ as an irrelevant negative control. Surface expression of pHLA complexes was measured using an anti-HLA class I antibody. Mean fluorescence Intensity (MFI) was used as a readout of HLA-peptide stability across a range of peptide concentrations. We observed varying degrees of HLA-B*58:01 surface stabilization by the tested peptides (**Figure 2A**). For IW9 and KW11, the magnitudes of responses were similar for the consensus and variant peptides tested, albeit less stabilizing than the positive control EBV peptide. In contrast, of the two TW10 variant peptide sequences detected in this cohort, both were associated with lower stability than the consensus HLA-B*58:01 peptide, which was close to the positive control EBV peptide.

**Figure 2:**
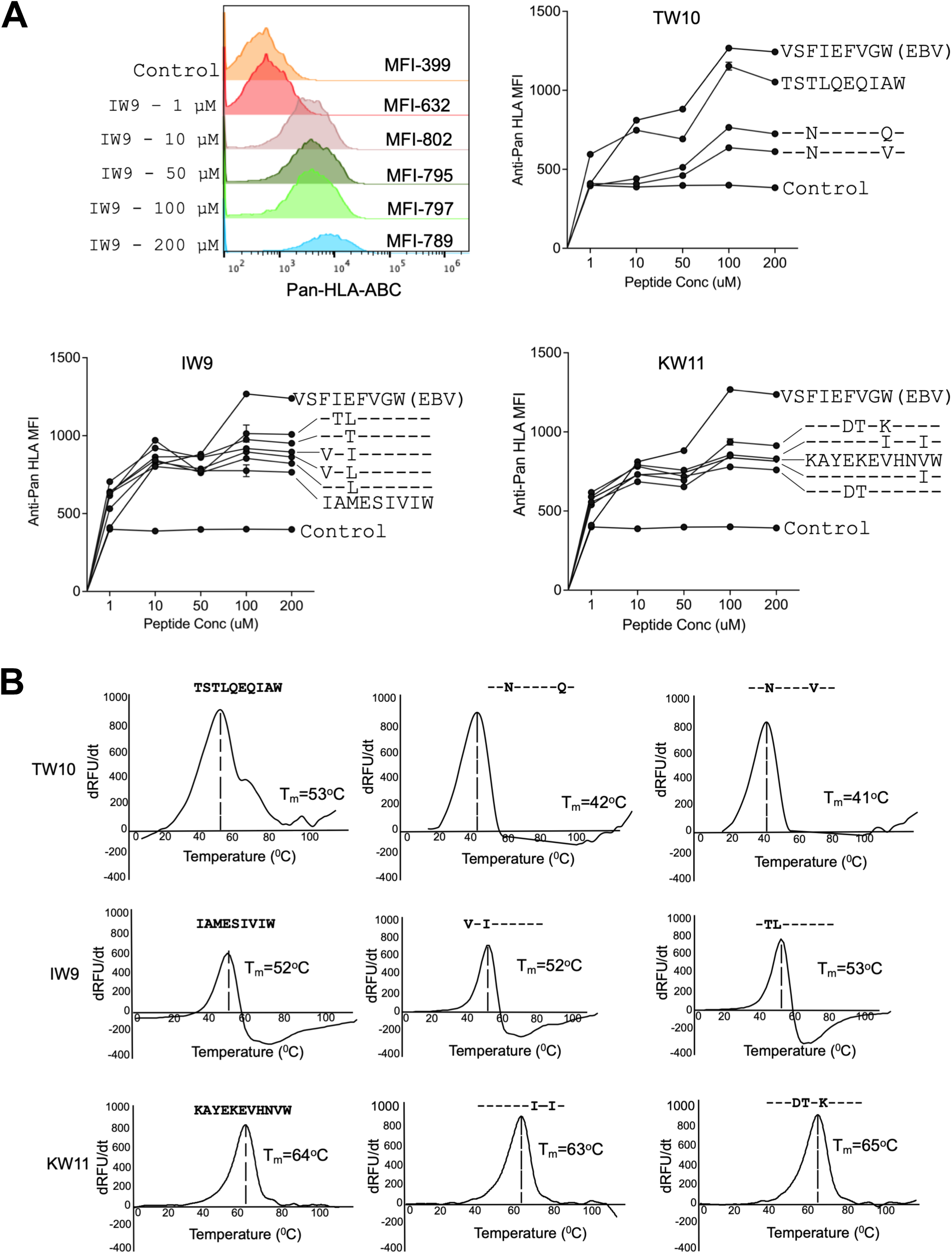
HLA-B*58:01 pHLA stability of consensus and variant epitopes. **(A)** Representative histograms for HLA mean fluorescence intensity (MFI) for consensus and variant peptides (concentration range 1-200 µM) using a TAP-deficient mono-allelic cell line engineered for constitutive expression of HLA-B*58:01. Shown are the MFI of pan-HLA staining. Each data point represents the mean of technical triplicates from experiments performed twice. **(B)** Relative pHLA stabilities by thermal denaturation profile of HLA-B*58:01 pHLA for the indicated HIV-1 consensus and variant epitopes. The x-axis depicts temperature. The y-axis depicts the derivative of temperature versus fluorescence (-dRFU/dT)^40^. The thermal stability (T_m_) for each HLA-B*58:01 pHLA complex is indicated.

To further define the ability of these peptides to be presented by HLA-B*58:01, we applied differential scanning fluorimetry (DSF)^40^. This assay measures the thermal stability of peptides bound to HLA as a function of temperature. Consistent with the HLA stability assay results, the variant peptides for IW9 and KW11 exhibited thermal stability similar to their respective consensus peptides. The TW10 results by DSF were consistent with findings of the HLA stability assays, with both TW10 variants showing reduced thermal stability compared to the consensus sequence (**Figure 2B**).

Collectively, these findings indicate heterogeneity in the ability of variant peptide-HLA-B*58:01 complexes to be presented for recognition. Among the three epitopes analyzed, KW11 peptides exhibited the highest thermal stability, indicating stronger peptide-MHC (pMHC) binding. Both the KW11 and IW9 variants were more efficiently presented compared to the TW10 variants. Notably, despite the overall stability of the three consensus peptides, multiple participants did not detectably target these epitopes in ELISPOT assays.

### Early ART preserves HIV-specific transitional memory CD8^+^ T cells

We next examined how antigen exposure duration influences the differentiation and transcriptional landscape of HIV-specific CD8^+^ T cells. Given the typically low frequency of pHLA tetramer^+^ T cells following ART-induced aviremia^26^, the strong stabilization of HLA-B*58:01 by the KW11 epitope and its variants, and because we detected *ex vivo* KW11-specific CD8^+^ T cell responses in multiple participants, we focused the analysis on this epitope. Peripheral blood mononuclear cells (PBMCs) obtained following prolonged aviremia on ART were enriched for CD8^+^ T cells and briefly treated with Dasatinib to preserve TCR-pHLA interactions^41^. The *ex vivo* KW11-specific CD8^+^ T cells were subsequently identified using both consensus and autologous variant pHLA tetramers and isolated by FACS. Sorted cells were subjected to scRNAseq using 10X Genomics technology. Resulting libraries underwent stringent quality control and processing as previously described^42,43^.

A total of 5,182 KW11-specific CD8^+^ T cells were obtained for downstream analysis (**Figure S2A and S2B**). Phenotypic profiling using CITE-seq revealed dominance of T stem cell memory (T_SCM_) cells, together with other subdominant memory subsets in both treatment groups (**Figure 3A**). Unsupervised transcriptomic clustering of these antigen-experienced tetramer-positive memory cells revealed 11 transcriptionally distinct clusters, spanning a differentiation spectrum from early memory (cluster 0) to NK-like cytotoxic effector memory (cluster 10) phenotypes. Modeling cluster composition using scCODA^44^ to compare KW11-specific CD8^+^ T cells from early versus late ART-treated donors revealed similarities and differences in the relative frequencies of the 11 identified clusters (**Supplementary material 1).** Notably, cluster 2 (C2), which was markedly enriched in persons with limited antigen exposure, exhibited gene expression features characteristic of transitional memory CD8^+^ T cells, including high expression of *CCR7, LEF1, CD62L* (SELL) and *TCF7*, (**Figure 3B**). This finding is consistent with prior studies showing that prompt initiation of ART during acute HIV infection preserves early-differentiated memory subsets^45–48^.

**Figure 3:**
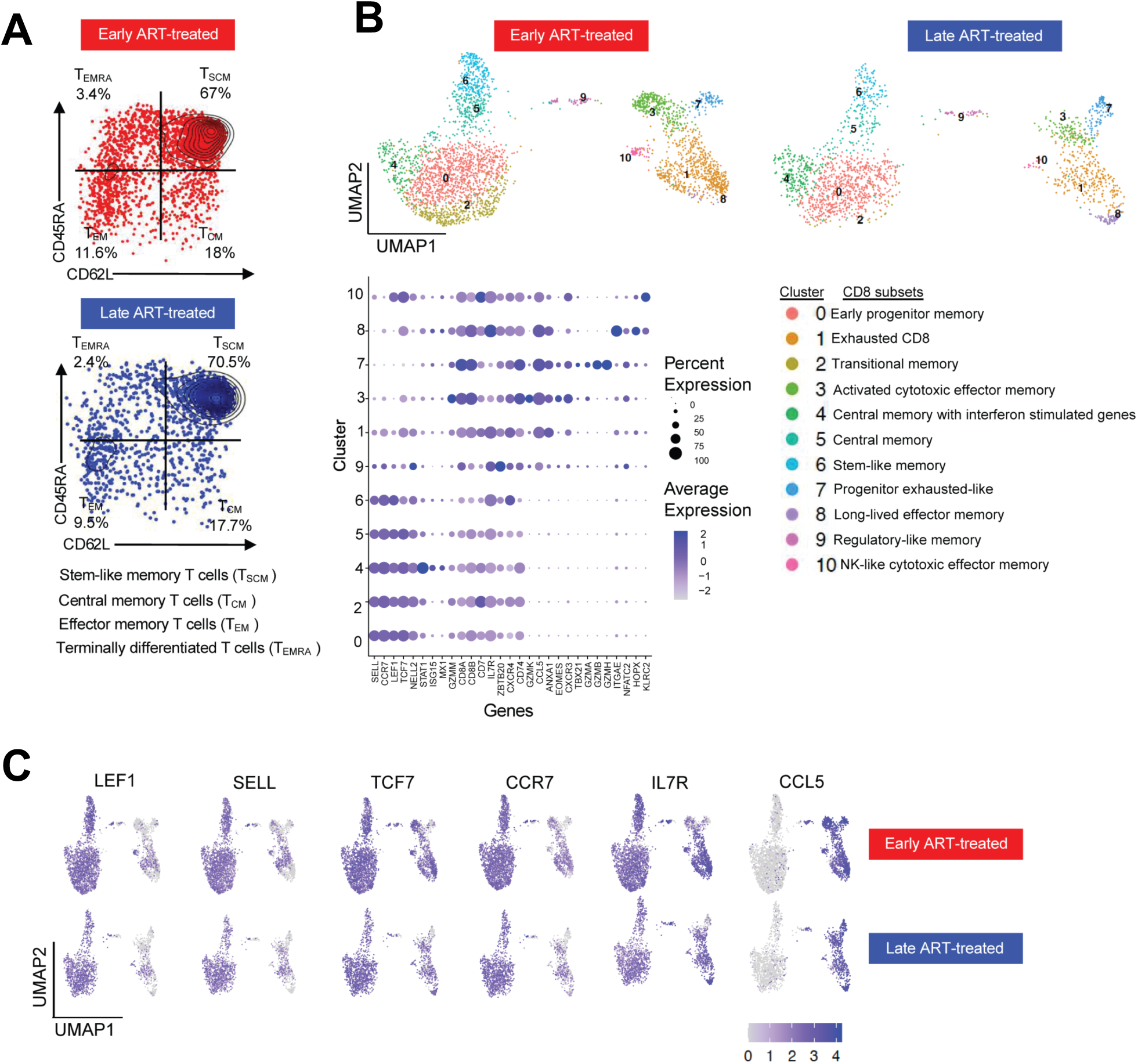
**Profile of *ex vivo* HIV-specific CD8^+^ T cells following ART treatment**: (**A**) Protein expression of CD45RA and CD62L from CITE-seq data illustrating the differentiation status of KW11-specific CD8^+^ T cells in early and late ART-treated donors. Quadrants correspond to distinct T cell subsets based on surface marker expression, quantified by unique molecular identifier (UMI) counts. (**B**) UMAP visualization of 5,182 KW11-specific CD8^+^ T cells from 12 donors (6 early and 6 late ART-treated). Cluster numbers are shown on the UMAP, with corresponding cluster names indicated. Transitional memory cells (cluster 2) were enriched in early ART-treated donors but depleted in the late-treated donors. The accompanying bubble size reflects the proportion of cells expressing each gene, while intensity (purple) indicates the average expression level. Clusters 0, 2, 4, 5, 6 and 9 correspond to memory-like populations, while clusters 1, 3, 7, 8 and 10 represent effector memory populations. (**C**) Log-normalized expression of selected genes defining central memory (left) and effector memory (right) clusters. For each gene, the top row represents clusters from early-treated donors, and the bottom row represents clusters from late-treated donors.

In contrast, clusters 1, 3, 7 and 10 represented transcriptionally distinct effector memory populations that were comparable in frequency in both early and late treated participants (**Figure 3C**). Cluster 8, annotated as long-lived effector memory CD8^+^ T cells, was notably enriched in the late ART-treated donor 272 (**Figure S2B**), who experienced the highest level and longest duration of viremia (see **Figure 1 and Table 1**) prior to ART initiation. These effector memory populations expressed genes linked to tissue residency and long-term persistence, such as CCL5, consistent with transcriptional programs observed in anti-tumor immunity^49^.

These data indicate that prolonged antigen exposure modestly remodels the CD8^+^ T cell differentiation landscape. The relative preservation of transitional memory populations in early ART-treated donors, contrasted with their loss following prolonged antigen exposure, reflects a transcriptional imprint of chronic antigenic stimulation.

### Prolonged viremia is associated with enhanced repertoire cross-reactivity

We next characterized the TCR repertoire recognizing HLA-B*58:01-stabilizing epitopes to determine how the duration of antigen exposure influences the clonal architecture and cross-reactivity profiles of individual TCR clonotypes. In order to focus on clonotypes with the ability to proliferate, as a measure of functionality^50^, and due to the small numbers of cryopreserved cells available, we stimulated PBMC with anti-CD3 and anti-CD28 to induce non-specific *in vitro* T cell expansion. We used barcoded pHLA tetramers constructed with the consensus and variant epitopes (**Table 3, Figures S3A-B**), followed by sorting for single-cell sequencing. Each tetramer was conjugated to a unique molecular identifier (UMI), enabling quantification of antigen-specific T cells at the single-cell level^51^. To facilitate multiplexed analysis, donor-derived CD8^+^ T cells (6,000-10,000 per sample) were barcoded using hashing antibodies^52^ and processed through the 10X Genomics platform^53^. CITE-seq and V(D)J libraries were first generated and sequenced using Illumina short-read technology. To validate these data and ensure high-confidence TCR assignments, the V(D)J libraries were further analyzed with Oxford Nanopore long-read sequencing. Only clonotypes detected by both sequencing platforms were considered validated and subsequently included in the final analysis. TCR clonotypes specific for peptide epitopes were detected with UMI counts ranging from as few as 1 to 160 (**Figure S4A-C**).

We assessed the capacity of individual TCR clonotypes, obtained following suppression of viremia by ART from early and late-treated individuals to recognize both the consensus HIV epitope and corresponding autologous variants. A clonotype was considered cross-reactive within an epitope family if it exhibited a UMI count ≥5 for a given consensus or variant peptide, ensuring specific TCR-pHLA interactions^53^. This threshold was selected to balance sensitivity and specificity while accounting for donor-specific hashing and minimizing noise from low-frequency events^53,54^. Using this criterion, we identified 79 unique KW11-specific clonotypes, 37 unique IW9-specific clonotypes, and 30 unique TW10-specific clonotypes in early-treated donors. In the late-treated donors, we detected 123 unique KW11-specific clonotypes, 34 unique IW9-specific clonotypes, and 24 unique TW10-specific clonotypes, all of which were included in downstream analyses.

For all 12 donors, clonotypes were present that targeted at least one of the epitopes tested. For all epitopes, TCR clonotypes that recognized variant epitopes and not the consensus sequence were detected (**Figure 4A, Figure S5A and S5B**). For example, the majority of KW11-specific clonotypes recognized variants and not the consensus sequence, and similar type-specific cells not targeting the consensus sequences were detected for IW9 and TW10 epitopes in smaller numbers (**Figure S5A and S5B**). Although cross-reactive TCRs capable of recognizing the consensus and all variant epitopes tested were also detected, these broadly cross-reactive clonotypes represented a small minority (<1%) (**Figure S5A and S5B**).

**Figure 4:**
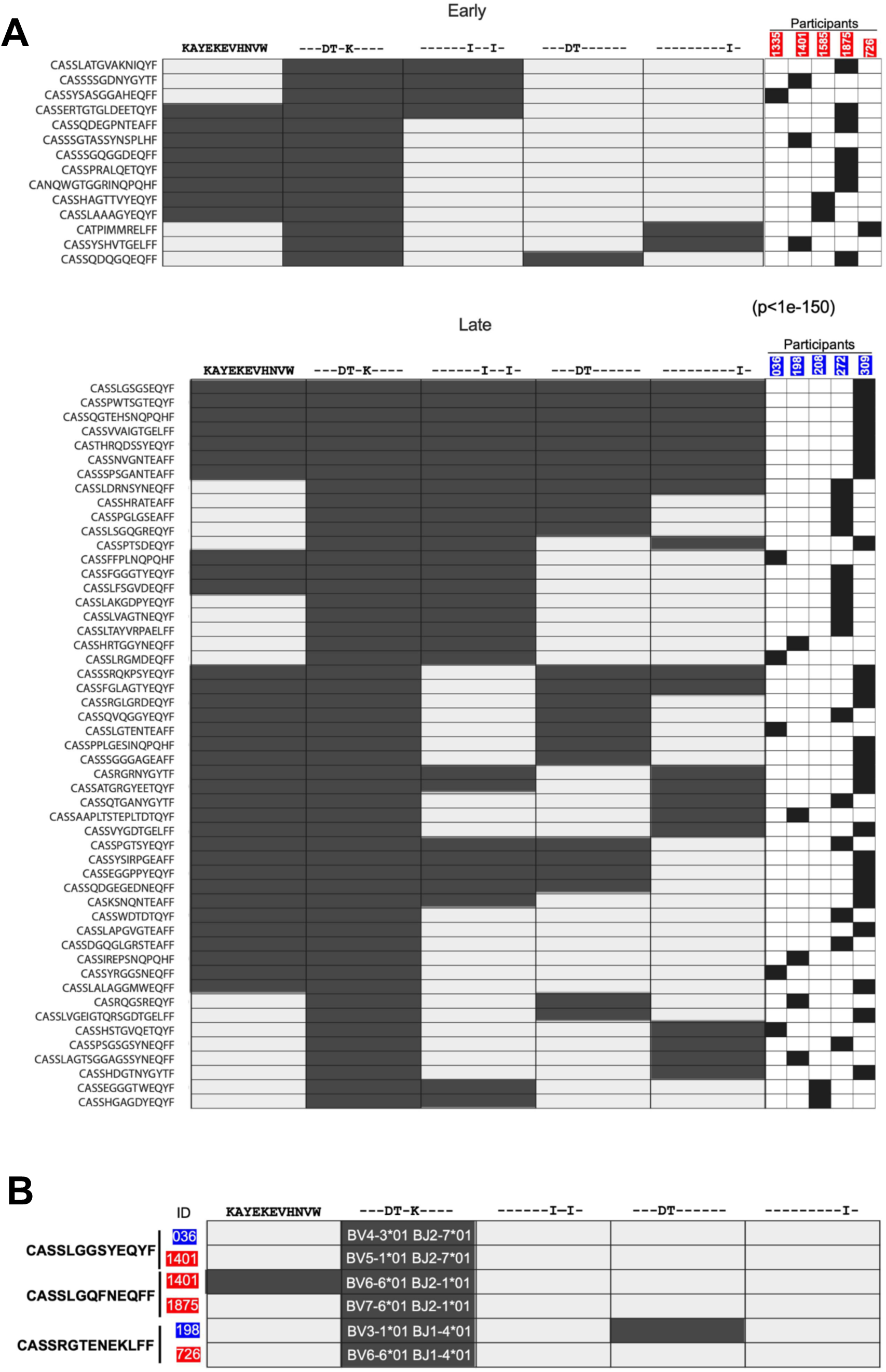
HIV-specific TCRs recognition in early and late treated groups. (**A**) Cross-reactive TCR clonotypes targeting the KW11 epitope variants in individuals who initiated ART during acute infection (top) versus those with delayed ART initiation (bottom), all assessed during sustained ART suppression. Each row represents a unique TCR clonotype, and each column indicates a KW11 epitope variant. Columns on the right show the donor-specific contribution of each clonotype included in the heatmap. Filled boxes indicate tetramer-positive clonotypes were obtained. 82% (65 of 79 total clonotypes) of KW11-specific clonotypes from early ART-treated individuals were monospecific (recognizing only one variant), compared with 52% 72 of 123in late ART-treated individuals. The non-cross-reactive clonotypes are not displayed in this heatmap. (**B**) Cross-reactivity profiles of the three identified KW11-specific public clonotypes. The shared CDR3β amino acid sequences are listed on the left, and the participant IDs are shown in red (early treated) and blue (late treated) boxes. Pairs that share a clonotype are indicated by the vertical lines. Recognition of variants is depicted by filled boxes. *TRBV* and *J* genes are indicated for each subject. To avoid cross-contamination or sampling error, each public clonotype was identified using a separate 10X cartridge. For each participant, the corresponding V and J genes are displayed in the second column from the left, aligned with the respective clonotype.

We next evaluated the impact of the duration of antigen exposure on the cross-reactivity of clonotypes targeting these peptide-HLA complexes, conducting a summary statistical analysis by randomly sampling clonotypes from the early- and late-treated groups for KW11, TW10 and IW9. Cross-reactive clonotypes recognizing at least two variants were more numerous in the late treated groups for KW11 (p<1e-150) and IW9 (p<0.0001). However, no significant difference was observed for TW10 (p=0.1973), a result that may be influenced by the limited number of clonotypes available for TW10.

These results indicate that prolonged antigen exposure drives an increase in the generation of HLA-B*58:01-restricted clonotypes specific for autologous variants. However, both limited and prolonged exposure to viremia lead to the generation of very few clonotypes able to recognize all peptide variants tested.

TCR sequencing also yielded CDR3 sequence data, providing us the opportunity to investigate the contribution of “public” TCR sequences, defined here as TCRs with identical CDR3β amino acid sequences found in at least two individuals^55–58^. We focused on TCR specific for KW11, for which we obtained the largest number of clonotypes. We identified three such public clonotypes out of the 202 clonotypes recovered (**Figure 4B**). Although these clonotypes shared identical CDR3β sequences, each was paired with a distinct Vβ gene segment. For instance, the clonotype CASSRGTENEKLFF was found in participants 198 (late-treated) and 726 (early-treated). While both used the same J gene (TRBJ-1-4*01), their V㊲ genes differed: TRBV3-1*01 in 198 and TRBV6-6*01 in 726. These findings indicate that identical CDR3β sequences targeting the same HIV variants can independently arise in different individuals under similar antigenic selection pressures, but that these appear to be rare in the context of HLA-B*58:01.

To assess whether differences in Vβ gene usage across individuals impact the recognition of epitope variants, we examined the reactivity of these public clonotypes against autologous KW11 variants. Notably, public clonotypes from donors 1401 and 198 exhibited cross-reactivity among KW11 variants, despite lacking shared Vβ genes (**Figure 4B**). In contrast, public clonotypes from participants 036, 1875, and 726 exhibited type-specific responses to the same KW11 variant. These data indicate that even among clonotypes with identical CDR3β sequences targeting identical pHLA complexes, differences in Vb usage exist, as do differences in cross-reactivity.

Overall, these results indicate that broadly cross-reactive HLA-B*5801-restricted clonotypes targeting all defined autologous epitope variants are inducible with either limited or prolonged viremia, but are not the dominant clonotypes. In addition, our results demonstrate that for these HLA-B*58:01-restricted epitopes, circulating variants are often more frequently targeted than the clade C consensus sequence.

### *TRBV* usage increases with chronic antigen stimulation

Given prior studies showing biased usage of certain Vbeta genes within the antigen-specific T cell repertoire^26,59^, we investigated the impact of antigen exposure duration on *TRBV* gene diversity among HLA-B*58:01-restricted responses (**Figure S6A-C**). Overall, shared *TRBV* genes were frequently observed between early and late-treated groups (**Figure 5A-C**). Notably, *TRBV5-1*01* and *TRBV4-1*01* were consistently enriched among HLA-B*58:01-restricted clonotypes, independent of the specific antigen recognized, suggesting that the interaction between HLA-B*58:01 and the TCR may bias *TRBV* gene selection.

**Figure 5:**
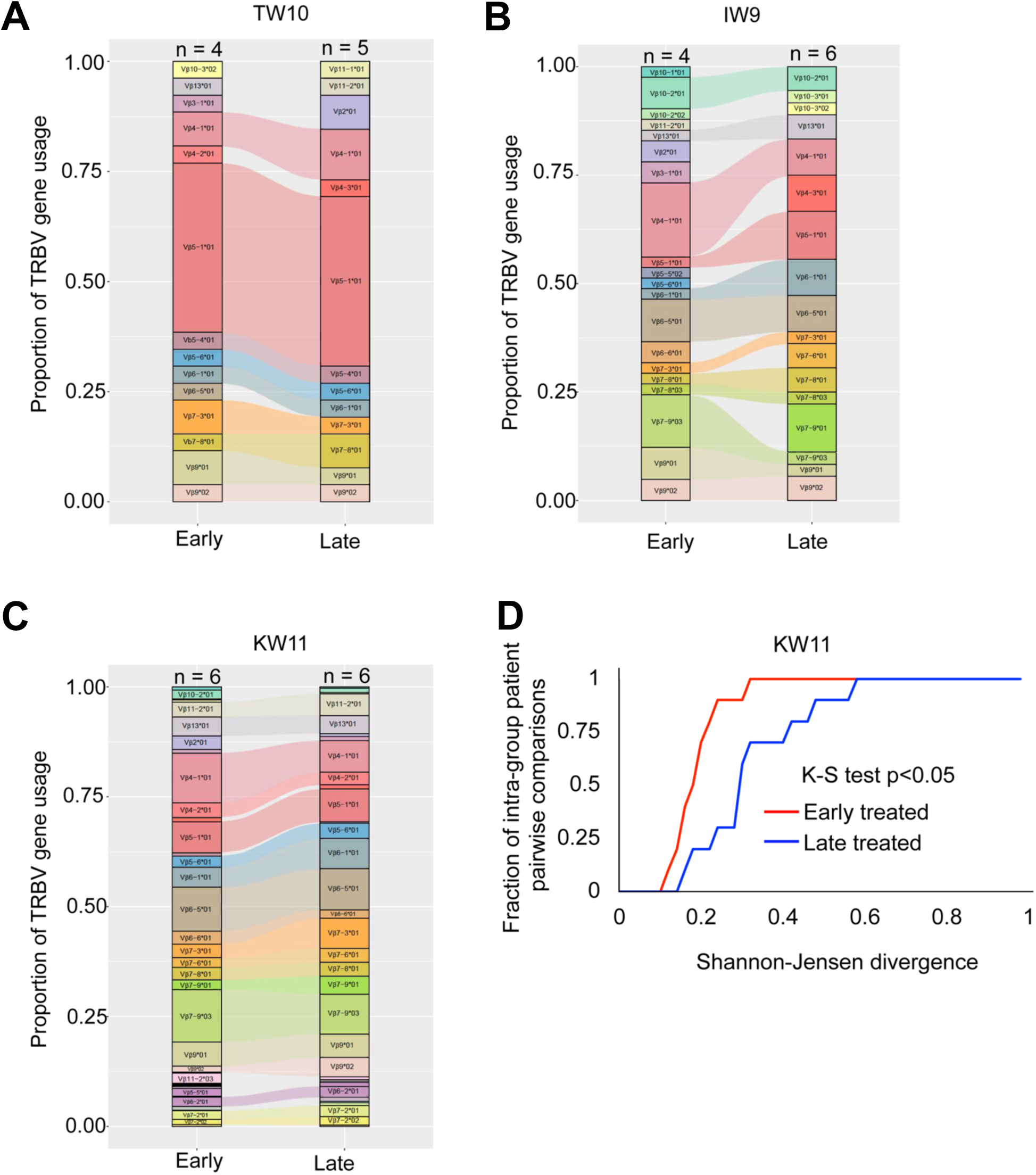
***TRBV* gene usage in HIV-specific clonotypes with varying antigen exposure durations**. (**A-C**) *TRBV* gene usage among TW10, IW9, and KW11-specific TCR clonotypes in early and late-treated participants. Each bar represents the frequency of shared *TRBV* genes within the epitope-specific repertoire, with distinct colors indicating different *TRBV* genes. The “n” above each bar denotes the number of donors contributing to the TRBV repertoire in each treatment group. (**D**) Cumulative distribution function (CDF) plot showing intra-participant comparisons for KW11 antigen, highlighting diversity in the TCR repertoire in response to different durations of antigenic stimulation. TCR diversity was assessed using Shannon-Jensen divergence (JSD)^61^.

To quantitatively evaluate TRBV diversity, we assessed pairwise divergence among early or late treated samples using the Shannon-Jensen (SJ) divergence metric, where lower values indicate reduced heterogeneity^60^. Due to the more limited number of clonotypes available for TW10 and IW9 antigens, we focused the SJ analysis to KW11-reactive clonotypes to ensure sufficient data for robust statistical comparisons within each group. Summarizing the cumulative distributions of pairwise SJ divergence within each group reveals that early-treated samples tend to display less TRBV divergence than late-treated samples (K-S test, KW11: p<0.05) (**Figure 5D**).

Although both groups shared multiple *TRBV* genes for HLA-B*58:01-restricted peptides, the cumulative distribution function plot for KW11 indicated that the limited antigen exposure group had more restricted *TRBV* gene usage, whereas the prolonged exposure group exhibited broader *TRBV* gene diversity.

### Prolonged viremia following acute HIV infection leads to enrichment in lower-avidity TCRs

To assess the impact of antigen exposure duration before ART initiation on the avidity of HIV-specific TCRs, we leveraged CITE-seq data to estimate the relative strength of TCR-pHLA interactions. Building on evidence that CD3 and TCR expression are directly correlated^62,63^, we quantified binding strength by calculating the ratio of barcoded pHLA tetramer UMIs to CD3 UMIs on a per-cell basis. To resolve differences in TCR affinity for cognate pHLA, cells were stained with a sub-saturating concentration of tetramer (1 nM for each specificity). Under these conditions, the extent of tetramer binding serves as a proxy for TCR strength, with higher-affinity TCRs binding more tetramers than their lower-affinity counterparts^64–69^. We quantified TCR binding strength by calculating the tetramer/CD3 UMI ratio for each clonotype and compared distributions between early and late ART-treated individuals. Collectively, early-treated individuals exhibited a significant enrichment of clonotypes with stronger tetramer binding across all three epitopes: KW11 (p<0.0001), IW9 (p=<0.0001) and TW10 (p=0.04) (non-parametric t-test; **Figure 6A**). In contrast, EBV-specific CD8^+^ T cells showed no difference in binding distributions between treatment groups, indicating that enhanced HIV-specific TCR sensitivity is driven by HIV antigen exposure history rather than generalized immune differences.

**Figure 6:**
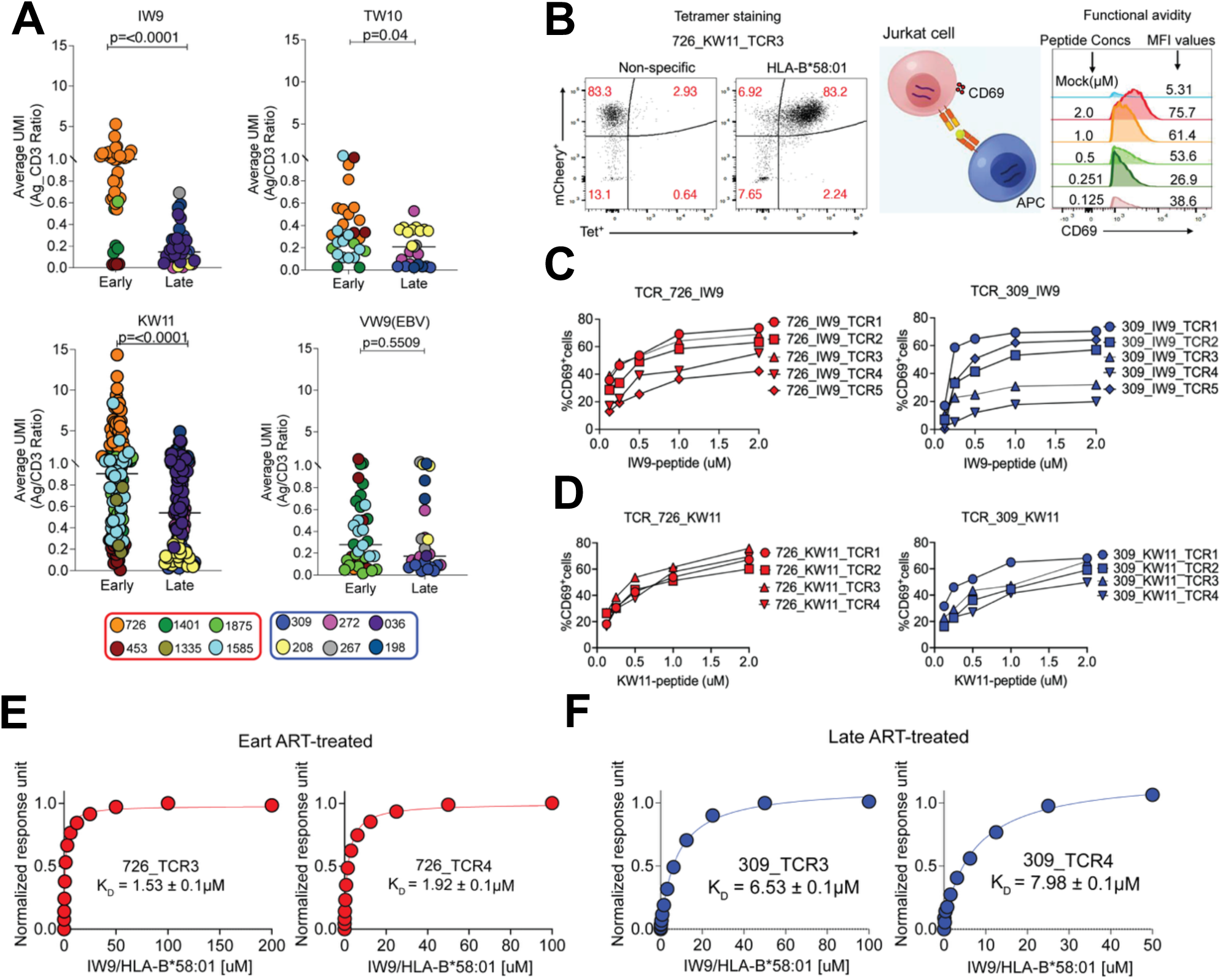
**Analysis of HIV-specific TCR avidity and functional affinity**. (**A**) Estimated TCR-pHLA binding strength for individual clonotypes, calculated as the average ratio of bound tetramer (antigen, Ag) UMIs to CD3 UMIs. Statistical comparisons were performed using a non-parametric t-test. Each donor contribution is depicted with a different colour matched to the donor. (**B**). Representative FACs plot showing a monoclonal Jurkat TCR stained with a non-specific control tetramer (B*42:01-TL9) and the cognate tetramer (B*58:01-KW11). TCR-engineered T cells were co-cultured with HLA-B*58:01 monoallelic antigen-presenting cells pulsed with graded peptide concentrations (2μM to 0.125μM) at a 1:1 ratio for 18 hours. The histograms illustrate dose-dependent CD69 expression for 726_KW11-specific TCR3 clone. (**C and D**). Representative peptide dose-response curves for (**C**) IW9-specific TCRs and (**D**) KW11-specific TCRs derived from early-treated donor 726 and late-treated donor 309. TCR identifiers correspond to clonotypes with paired α and β-chains confirmed by both Illumina and Oxford nanopore sequencing. (**E and F**) Representative steady-state SPR measurements showing binding interactions between soluble HLA-B*58:01-IW9 pHLA complex and four IW9-specific TCRs, two from the early ART-treated donor 726 (**E**) and two from the late ART-treated donor 309 (**F**). Individual plots indicate donor TCR IDs and corresponding dissociation constants (K_D_)

Importantly, the distributions of tetramer/CD3 UMI ratio were strongly donor-specific, a pattern observed for both HIV- and EBV-specific CD8^+^ T cells. This finding is consistent with prior evidence from both mouse and human studies demonstrating individual variability in TCR/CD3 expression and clonotype-intrinsic properties^70,71^. Thus, these absolute ratio values should be interpreted with caution, taking donor effects into account. Emphasis should instead be placed on within-epitope, between-group comparisons, where meaningful biological differences are more reliably detected, as evidenced in this study.

To functionally assess these TCRs, we selected 42 IW9 and KW11-specific clonotypes spanning a range of tetramer/CD3 ratios from two early and two late-ART-treated donors. We successfully re-expressed 35/42 (83%) TCRs in Jurkat reporter cells. Antigen sensitivity was quantified by stimulating TCR-transduced cells with peptide-pulsed TAP-deficient, mono-allelic HLA-B*58:01*-* expressing cells and measuring CD69 upregulation as a readout of activation^72–74^. Dose-response curves were generated and EC_50_ Values(-log_10_) were estimated for each TCR (**Figure 6B**).

All TCRs tested retained functional responsiveness across the peptide titration range (**Figure 6C-D and Figure S7**). Notably, tetramer/CD3 ratios did not correlate with functional EC_50_ values (**Figure S8**), indicating that tetramer binding strength and functional antigen sensitivity represent distinct physiological parameters that should be independently evaluated when characterizing TCR avidity.

To directly compare TCR affinities between early and late ART-treated donors, we performed surface plasmon resonance (SPR) on soluble IW9-specific TCRs generated from donors 726 (early treated) and 309 (late treated). Four TCRs were successfully expressed and assessed for binding to the IW9-HLA-B*58:01 complex. TCRs from the early-treated donor bound with substantially higher affinity (726_TCR3: K_D_ = 1.53 ± 0.1µM; 726_TCR4: 1.92 ± 0.1µM) (**Figure 6E**), whereas those from the late-treated donor showed markedly weaker binding (309_TCR3: K_D_ = 6.53 ± 0.1µM; 309_TCR4: K_D_ = 7.98 ± 0.1µM) (**Figure 6F**).

Taken together, these findings indicate that prolonged antigen exposure prior to ART initiation skews the HIV-specific TCR repertoire toward lower-avidity clonotypes, thereby reducing T cell responsiveness to antigen. In contrast, early ART preserves a pool of higher-avidity TCRs, with enhanced functional sensitivity.

## Discussion

In this study, we investigated the impact of antigen exposure duration on the clonal diversity, functional quality, and differentiation phenotype of the virus-specific CD8^+^ TCR repertoire, using acute HIV-1 infection as a model. We leveraged a unique cohort of PWH identified during hyperacute infection, all matched for the same HLA class I allele and tested for recognition of the same HIV epitopes restricted by this allele. We observed that ART-induced limitation of viremia was associated with the enrichment of relatively higher-avidity HIV-specific TCRs, in contrast to the lower-avidity TCRs and relative loss of transitional memory CD8^+^ T cells observed following prolonged viremia. Notably, cross-reactive TCRs capable of recognizing circulating viral variants were detectable in both early and late ART-treated groups, and were broader in those with prolonged exposure for two of the three epitopes evaluated. However, only a limited subset exhibited broad recognition across all autologous variants within the antigen family.

Studies of human cellular immune responses have been limited by HLA diversity and the inability to modulate antigen exposure. Here, we utilized samples from the FRESH cohort^25,28^, in which sufficient numbers of persons with hyperacute infection were identified to allow us to match all participants for a dominant HLA class I allele, as well as to stratify by duration of exposure to viral antigen due to timing of ART administration. Using barcoded tetramers constructed with these peptide epitopes and their variants, we identified clonotypes capable of recognizing all circulating variants, which is critical for controlling a rapidly evolving virus like HIV, and likely critical for T cell vaccine development. However, despite being inducible, these broadly cross-reactive clonotypes were rare and did not dominate the TCR repertoire in natural infection.

Using this same panel of barcoded tetramers, we also examined the avidity of the responses induced by both immediate and delayed administration of ART. At the clonotypic level, we showed that immediate ART results in the enrichment of higher avidity TCRs, compared to lower avidity responses seen with delayed ART. Although previous studies have shown that TCR engagement can lead to surface TCR downregulation^75^, in our experimental approach, expanded cells were rested to allow TCR re-expression before tetramer staining. We then used the tetramer-to-CD3 ratio as a quantitative metric of avidity and validated representative TCRs spanning a range from high to low avidity. Importantly, no differences were observed in the avidity of contemporaneous control EBV-specific TCRs between the two groups, indicating that the observed differences in avidity of HIV-specific clonotypes among these participants were specific to HIV antigens. Our findings align with the recent studies describing the evolution of dominant clonotypes from high to low avidity in chronic CMV infection^24^ and in cancer model systems^76^. Persistent antigen stimulation can result in the apoptosis or cellular senescence of high-avidity T cells that are initially selected during acute infection^68,77^. Over time, this process may drive a shift in clonal dominance from high to low avidity T cells, reflecting the adaptive dynamics of the T cell repertoire in response to prolonged antigenic challenges^78,79^.

Consistent with this model, we observed a relative depletion of transitional memory CD8^+^ T cells following prolonged viremia before ART initiation. The mechanism underlying this selective loss remains unclear, but one possibility is that this subset is susceptible to activation-induced exhaustion and death during chronic stimulation. Thus, while memory differentiation trajectories appear to be shaped primarily at the transcriptional level, *TRBV* usage within this subset likely reflects an intrinsic repertoire constraint.

Our findings have important implications for the design and interpretation of HIV analytical treatment interruption (ATI) studies. CD8^+^ T cell functionality has been associated with the ability of rare individuals to achieve post-treatment control following treatment interruption^80–83^. Here, we show that individuals who initiated ART during acute infection retain more avid HIV-specific TCRs compared with those treated during chronic infection. This enrichment provides a potential mechanistic explanation for improved ATI outcomes in early-treated individuals^84,85^. High-avidity TCRs would be expected to detect lower levels of antigen presentation, allowing CD8^+^ T cells to identify and eliminate infected cells at the earliest stages of viral reactivation as well as low level antigen presentation that may be present in cells comprising the HIV reservoir^86^. Such heightened sensitivity may facilitate more effective and durable immune containment of recrudescent virus. Conversely, late-treated individuals, who harbour a greater proportion of low-avidity clonotypes, may have diminished capacity to sense low-density antigen, potentially impairing CD8^+^ T cell engagement and reducing the likelihood of sustained control during ATI.

Together, these results indicate that early ART initiation preserves a functionally superior HIV-specific TCR repertoire that may confer a measurable advantage during ATI. This aligns with recent ATI studies demonstrating enhanced viral control among early-treated cohorts^84,85,87^ and highlights TCR avidity as a potential biomarker for predicting ATI outcomes.

This study has certain limitations worth noting. First, we focused exclusively on CD8^+^ T cell responses restricted by the *HLA-B*58:01* allele, an allele associated with better outcomes in HIV infection. While informative, this approach does not capture the diversity of HLA-restricted responses across other alleles that may also contribute to HIV control. Second, our functional assessments relied on *ex vivo* and *in vitro* assays, which, although powerful, may not fully reflect the complexity of TCR responses *in vivo*, where interactions with other immune cells and tissue-specific factors shape responsiveness. Similarly, the use of TCR-transduced Jurkat cell lines to measure antigen sensitivity provides a controlled system but may not fully recapitulate sensitivity on infected target cells^20^, even when validated on peptide-loaded APCs. To address this, we complemented these assays with SPR to validate representative TCRs. Third, although we examined cross-reactivity against autologous viral variants identified during the acute phase of infection, the evolving nature of HIV poses challenges, as viral escape can limit the long-term relevance of cross-reactive clonotypes identified at a single time point. Finally, the relatively small cohort size, the focus on epitopes presented by a single class I HLA allele, and the study of repertoires induced by only a few epitopes, albeit studied in depth, indicate that broader studies will be helpful in addressing generalizability. Future studies incorporating longitudinal analysis and larger cohorts, along with multi-dimensional assessments of immune responses across additional HLA alleles, would be valuable in providing a more comprehensive understanding of TCR dynamics and their role in HIV control.

In summary, this study underscores the influence of antigen exposure duration in shaping the HIV-specific T cell repertoire in terms of cross-reactivity, avidity and differentiation of memory subsets. Moreover, by focusing on HLA-B*5801 restricted epitopes, the data show dimensions of differing antiviral pressure mediated through an allele that is associated with better outcomes in some but not all persons expressing this restriction element. Despite focusing on a single HLA allele and a relatively small sample size, the use of advanced techniques such as barcoded tetramers and single-cell TCR sequencing provided a high-resolution analysis of clonotypic diversity, functional quality, and lineage relationships. These findings highlight the potential of early ART initiation to promote higher-avidity TCRs and preservation of stem-like memory cells. These insights underscore TCR avidity as a promising biomarker for predicting outcomes during ATI and for identifying individuals most likely to benefit from personalized ATI strategies. More broadly, leveraging insights into TCR avidity and memory-state preservation may inform the development of next-generation immunotherapies for HIV and other chronic viral infections.

## Materials and Methods

### Study design

This study utilized cryopreserved peripheral blood mononuclear cells (PBMCs) obtained from the FRESH cohort participants. HLA typing of these samples was performed at Dr. Mary Carrington’s laboratory at the National Cancer Institute in Fredrick, USA, following previously described methods^32^. The twelve participants included in this study were preselected based on the availability of PBMC samples and the detection of HLA-B*58:01 restricted responses by IFN-g Elispot during the acute phase of HIV-1 infection. The cohort size was determined based on the availability of sufficient peripheral blood mononuclear cells (PBMCs) rather than a pre-specified effect size. Each participant provided written informed consent in accordance with protocol no. BF298/14, approved by the Biomedical Research Ethics Committee of the University of KwaZulu-Natal, and protocol no. 201P001018/PHS, approved by the Institutional Review Board of Massachusetts General Hospital and the Partners Human Research Committee.

### ELISPOT assays

IFN-gamma ELISPOT assays were performed according to the manufacturer’s instructions with the human IFN-gamma ELISPOT Basic Kit (ALP) (Cat. 3420-2 A, Mabtech). Briefly, 2 X 10^5^ peripheral mononuclear cells (PBMCs) were seeded in individual wells of the IFN-gamma-coated plate with each peptide at a final concentration of 1 μg/ml and incubated overnight. 0.2 μM PHA (Cat. 11249738001, Sigma) and 1 μM CMV peptides (Cat. 3619-1, Mabtech) were used as positive controls. An equivalent amount of RPMI-1640 medium with DMSO (Cat. 317272, EMD Millipore) was used as a negative control. Responses were considered positive if they exhibited at least 3 times the mean number of spot-forming units (SFU) observed in the three negative control wells, and a minimum of 50 SFU/10^6^ PBMCs.

### Viral sequencing

Viral sequencing was carried out following established protocols^88^. Briefly, DNA extraction was conducted using QIAGEN DNase Blood and Tissue Kits (QIAGEN). Total HIV-1 DNA and host cell DNA concentrations were estimated using Bio-Rad ddPCR, with primers and probes targeting Gag, Env, Nef, and Pol protein regions. Extracted DNA was quantified via ddPCR, and subsequent dilutions were made based on these results. Full-length viral sequences were obtained using a single-amplicon nested PCR approach for HIV-1 full genome amplification. All PCR amplicons were prepared for sequencing using MiSeq (Illumina) library preparation through shearing-based adaptor-ligation technology, followed by MiSeq sequencing. The resulting sequencing data were analyzed as previously described^88^.

### Peptide synthesis and analysis

All peptides were synthesized by the Mass General Peptide Synthesis Core following established protocols^89^.

### HLA-B*58:01 expressing cell line

The HLA-B*58:01-expressing cell line was kindly provided by Dr. Gaurav Gaiha (Ragon Institute of MGH, MIT, and Harvard). A TAP-deficient, mono-allelic HLA-B*58:01 expressing cell line was generated through sequential transduction of the HLA-null human B cell line 721.221, following a previously described protocol^37^.

### HLA-B*58:01 peptide stability assay

To assess peptide stability, fifty thousand TAP-deficient monoallelic HLA-B*58:01-expressing 721.221 cells were incubated with varying concentrations of peptides (1-200 μM) and 3 μg/ml of β2M (Sigma-Aldrich; Sino Biological) in RPMI-1640 medium overnight at 26 °C with 5% CO_2_ for 18 hours. Parallel controls, without peptide but with equivalent concentrations of DMSO, were included. Following incubation, cells were shifted to 37 °C with 5% CO_2_ for 3 hours and subsequently stained for viability and pHLA class I surface expression using HLA-ABC (W6/32) APC (1:100 Biolegend).

### Construction and expression of soluble major histocompatibility complex (HLA) proteins

The HLA-B*58:01 pHLA complex was expressed following a previously described protocol^90^. DNA sequences encoding the ectodomains of the HLA-B*58:01 heavy chain were optimized for expression in the *E. coli* system and subsequently cloned into the pET22b(+) vector (Cat. 69744-3, Novagen) between the NdeI (Cat. R0111S, NEB) and Xhol (Cat. R0146S, NEB) restriction enzyme sites. The transformed *E. coli* were induced for inclusion body expression by adding 1 mM IPTG (Cat. 156000, RPI) to the culture at an OD:600. The cells were incubated for 4 hours at 37°C, after which the cell pellets were collected, resuspended, and lysed in extraction buffer 50 mM Tris-HCl (Cat. 46-031-CM, Corning), 100 mM NaCl (Cat. 71382, Sigma), 2% Triton X-100 (Cat. A16046.AP), pH 8.2 containing fresh lysozyme (Cat. L6876, Sigma), DNase-I (Cat. 10104159001, Sigma), and PMSF (cat. 36978 ThermoFisher). The lysates were sonicated, and the inclusion bodies were collected by centrifugation at 10,000 rpm for 10 minutes. To ensure purity, the inclusion bodies were resuspended and sonicated again, followed by three washes with wash buffer (50 mM Tris, 20 mM EDTA, pH 8.0). β2m (light chain) was expressed and purified following the same protocol. The constructs for both β_2_m and the HLA class I heavy chain were generously donated by Dr. Xianlong Li (University of Science and Technology of China, Hefei).

### Refolding and purification of peptide-HLA

All the consensus and variant peptides used for the thermomelt assay were synthesized by the MGH peptide synthesis core facility. For the refolding process, heavy chain (56 mg) and β_2_m (10 mg) were diluted in 1 liter of a refolding buffer containing 100 mM Tris-HCl (pH 8.0), 0.4 M arginine (Cat. A5131, Sigma), 0.5 mM oxidized glutathione (Cat. G4376, Sigma), 1.5 mM reduced glutathione (Cat. G4251, Sigma), 2 mM EDTA (Cat. E4884, Sigma), 4 M urea (Cat. U5128, Sigma), and 0.2 mM PMSF. The refolding reaction was performed in a total volume of 1 liter over 72 hours at 4 °C. Following refolding, the solution was dialyzed multiple times against 10 mM Tris-HCl (pH 8.0) at 4 °C using a 6-8 kDa molecular weight cut-off dialysis membrane (Cat. 132670, Spectrum). The dialyzed solution was then concentrated and subjected to size-exclusion chromatography using a Superdex 75 (GE Healthcare) gel filtration column with a running buffer of 10 mM Tris-HCl (pH 8.0) to remove any misfolded proteins. Finally, a Superdex 200 increase column (GE Healthcare) was used for buffer exchange. Protein concentrations were determined using a NanoDrop spectrophotometer

### Differential scanning fluorimetry

Differential scanning fluorimetry was performed using the Bio-Rad CFX-96 Real-Time PCR system. For thermal stability measurements, 20 μL of 2 μM pHLA was mixed with 0.2 μL of 100X SYPRO orange dye (Cat. 4461146, ThermoFisher). The temperature was increased at a rate of 1 ^0^C/min, ranging from 20 °C to 95 °C. Data analysis was performed using OriginPro, following the methodology described by Hellman et al.^91^. The apparent melting temperature (T_m_) was determined by identifying the point where the fluorescence signal indicated a 50% transition.

### Generation of barcoded tetramers

Barcoded tetramers for HLA-B*58:01 epitopes were generated using HLA-B*58:01 easYmer obtained from ImmunAware (Horsholm, Denmark). The easYmer is a highly active formulation of a peptide-receptive heavy chain HLA-B*58:01 molecule complexed with the light chain *β*_2_-microglobulin (*β*_2_m) molecule and di-peptide, which is highly sensitive to subtle temperature changes. A peptide exchange assay was performed by incubating easYmer with the peptides of interest at 18 °C for 48 hours to generate monomers, which were then used to create tetramers. Each peptide-HLA tetramer was uniquely DNA barcoded using BioLegend’s TotalSeq C streptavidin. These tetramers were used to stain PBMCs from participants.

### Non-specific T cell expansion

PBMCs were stimulated with soluble anti-CD3/CD28 (CD3: Mabtech, Cat. No: 3605-15; CD28: BioLegend, Cat. No: 302934) using 2 μg/mL anti-CD3 (OKT3) and 1 μg/mL anti-CD28 (CD28.2) for 6 days, followed by a 2-day resting period before tetramer staining of the expanded samples.

### Tetramer staining and FACS cell sorting

Expanded cells were initially incubated with 0.04% dextran sulfate and 0.3% of monocyte blocker (Biolegend, Cat. No: 426102) for 10 minutes at 4 ^0^C. The cells were then stained with a cocktail of sixteen unique barcoded tetramers at a 1nM final staining concentration for each tetramer for 30 minutes at 4 °C. Following staining, the cells were washed three times for five minutes at 300g. Subsequently, they were incubated with an FcX blocker (Biolegend, Cat. No: 422301) and then stained with both CITE-seq and FACS antibodies (details in the table below). Each sample was labeled using a unique β_2_m-based hashing antibody (BioLegend, Cat. No: 394633-65). Flow cytometry and cell sorting were performed at the Ragon Institute Flow Cytometry and Imaging Core using a FACSAria sorter. Flow cytometric analyses were performed using FlowJo v10.0.9 (TreeStar).

### *Ex vivo* single-cell RNA-seq processing

Single-cell gene expression, surface protein (CITE-seq), and TCR libraries were processed simultaneously using CellRanger v7.0.1 in multi-mode against the GRCh38-2020-A reference and vdj_GRCh38_alts_ensembl-5.0.0 annotations. Gene and protein count matrices were imported into R v4.2.0 and analyzed using Seurat v4.1.0.9007. Extensive filtering was applied based on hashing antibody and tetramer UMI counts: Cells were retained only if they contained a minimum of 1,000 UMIs for the assigned hashing antibody and less than 200 UMIs for all other hashtags (to confirm sample identity). Antigen specificity was assigned by requiring ≥5 UMIs for a given pHLA tetramer, with no detectable signal from tetramers outside the corresponding epitope family. Barcodes corresponding to TCRβ sequences independently identified in the expanded dataset were added if missing due to filtering, primarily affecting KW11-reactive cells from Participant 272. Following filtering, cells contained 518–12,208 transcripts (median 3,270) and 333–3,857 detected genes (median 1,336). TCRβ V(D)J gene calls as well as CDR3 nucleotide and amino-acid sequences were retrieved from the CellRanger AIRR outputs and incorporated into the Seurat object as metadata.

The dataset was log-normalized to 10,000 transcripts per cell using Seurat’s NormalizeData function. Variable genes (n = 2,000) were identified using FindVariableFeatures with the variance-stabilizing transformation. Data were scaled and centered with scale.max = 10, block.size = 1,000, and min.cells.to.block = 3,000.

Principal component analysis (PCA) was performed with 30 PCs, which were used for downstream analysis. UMAP embeddings were generated using RunUMAP with the following parameters: n.neighbors = 30, method = “uwot”, metric = “cosine”, learning.rate = 1, min.dist = 0.3, spread = 1, set.op.mix.ratio = 1, local.connectivity = 1, repulsion.strength = 1, negative.sample.rate = 5, uwot.sgd = FALSE, seed.use = 42. For clustering, a kNN graph (k = 20) was constructed using the first 30 PCs (RANN, nn.eps = 0, Euclidean distance). Louvain-based SNN modularity optimization was performed at resolution 0.8 (10 starts, 10 iterations), yielding 11 transcriptionally distinct clusters. Marker genes for each cluster were identified using FindAllMarkers (Wilcoxon test) and visualized using dot plots. Cluster identities were assigned based on recent published CD8⁺ T-cell differentiation markers described by Masopust *et al*.,^92^ and Galletti *et al*.,^47^. Differential gene expression between early- and late-ART cells was assessed using a Wilcoxon rank-sum test within each cluster.

Cluster frequency differences between early and late ART groups were evaluated using scCODA^44^. Cluster-level cell counts for each sample were tabulated and analyzed in a Python 3.11.6 environment, using cluster 9 (regulatory-like memory cells) as the reference category due to its stable representation across samples. Model covariates included: treatment timing (early versus late ART), donor of origin, and donor x time interaction. Compositions were estimated using Hamiltonian Monte Carlo sampling. Effect sizes were summarized at multiple credible intervals (False discovery rate (FDR) thresholds: 0.05, 0.1, 0.2, 0.3, 0.4) and reported in Supplementary material 1. Results were considered significant if the final parameter estimate excluded zero, and effect sizes were expressed as log2-fold changes relative to the reference cluster.

### Single-cell TCR long read sequencing

Paired participant samples, (early and late-treated individuals) were processed on a 10X Chromium X instrument, generating cDNA using standard 5’ DGE RT chemistry and 12 cycles of PCR with Feature cDNA primer4, then split into a polyA+ cDNA and a cDNA Feature barcode fraction (SPRI size selection). The polyA+ cDNA was used to generate gene expression (GEX) and TCR VDJ libraries. The quality and quantity of TCR libraries were evaluated using an Agilent Bioanalyzer High Sensitivity DNA kit. GEX and Feature Barcode libraries were sequenced on an Illumina NextSeq 500 instrument in paired-end mode (26nt/90nt), with 10% of the reads allocated to FB and GEX and the rest to TCR libraries. VDJ libraries were prepared with Oxford Nanopore Technology (ONT)’ SQK-NBD114 kit, with barcodes 1 to 7 to identify 10X libraries of origin and sequenced on an ONT Promethion instrument, using a FLO-PRO114M flowcell for 50 hours, in high-accuracy 400 bps base calling mode.

### Sequencing data processing

Post-sequencing, Illumina FB data were processed using 10X’s CellRanger v.7.0.1 on the cloud and as a command-line version on a compute cluster, using CellRanger’s GRCh38-2020-A reference. Cell barcodes and unique molecular identifiers (UMIs) were retrieved from long read data using the epi2me/wf-single-cell v.0.2.3 pipeline, mapping the cleaned reads against the GRCh38-2020-A reference, running nextflow 23.0.1 in a Singularity environment [https://github.com/epi2me-labs/wf-single-cell]. Reads mapping to the TCR alpha and beta loci on chromosomes 14 and 7, respectively, were retrieved and split based on their cell barcodes into fasta-formatted files. These files were downsampled to 500 reads per cell barcode, and contigs were assembled using canu, followed by three rounds of polishing with racon and remapping using minimap2, as adapted from^93^, followed by a final round of polishing with ONT’s medaka. TCR V, D, and J usage as well as CDR3 sequences and junctions were called using a standalone version of IgBLAST (v.1.21.0) against IMGT V-Quest 202312-3’s collection of human TCR gene segments^94,95^. Cell barcodes with unique productive TCR alpha or beta calls were considered first. For cell barcodes with more than one productive TCR beta or two TCR alpha chains, collapsing of the calls at the VDJ and CDR3 amino-acid levels was performed, and only cell barcodes meeting the unique TCR beta and/or two TCR alpha criteria were kept.

The resulting sequence data were intersected with the 10X FB data by exact matching of cell barcodes, and the data were post-processed using Seurat v.3 in the R 4.2.0 statistical environment^96^. Patient-specific stringent filtering criteria were applied to retain only cells with convincing tetramer staining by excluding cells with UMI signals on two distinct antigens, which were submitted to repertoire analysis using the vdjtools software package [https://vdjtools-doc.readthedocs.io/].

Shannon-Jensen divergence of TCRBV gene usage was calculated for all donor pairs within early- or late-treated groups, and the significance of differences among their cumulative distributions was assessed using the Kolmogorov-Smirnov (KS) test (R statistical environment). VDJ gene usages were also visualized as alluvial plots using the ggplots and ggalluvial libraries. For the purpose of estimating TCR-HLA/peptide avidity, log10-transformed ratios of antigen-specific tetramer counts over CD3 CITE-Seq counts (with a pseudo count of 1) were computed.

Sequencing relationships among CDR3 junctions were assessed for each antigen-binding subgroup. Briefly, unique CDR3 junctions of lengths 11-16 amino acids were submitted for multiple sequence alignment using the combination of multiple sequence aligners, as implemented in M-Coffee (T-Coffee online alignment server, https://tcoffee.crg.eu/)^97^. Circular Newick phylogenetic trees recapitulating sequence relationships were visualized using the online iTOL server (https://itol.embl.de/)^98,99^.

### Soluble TCR Expression and Purification

Recombinant soluble TCRs used for SPR experiments were produced using baculovirus expression in Hi5 insect cells, as previously described^100^. Briefly, the extracellular domain sequences of the TCR alpha chain were cloned into the pAcGP67a vector, which included a 3C protease site (LEVLFQGP), an acidic leucine zipper, an AviTag biotinylation site (GLNDIFEAQKIEWHE), and a poly-histidine tag (6xHis). Similarly, the extracellular domain sequences of the TCR beta chain were cloned into the pAcGP67a vector, containing a 3C protease site followed by a basic leucine zipper and a poly-histidine tag. For expression, 2 mg of each plasmid was transfected into SF9 insect cells along with BestBac 2.0 linearized baculovirus DNA (Expression System) using Cellfectin II reagent (Invitrogen). The primary transfection supernatant was then used to infect SF9 cells at a 1:1000 dilution to amplify the virus. Titration of the alpha and beta chain viruses was performed to determine the optimal ratio for efficient heterodimer formation. Protein production was scaled up by infecting Hi5 cells with the optimized virus ratio. The secreted proteins were then purified using HisPur Ni-NTA resin (Thermo) via immobilized metal affinity chromatography (IMAC).

### Surface plasmon resonance experiments

Surface plasmon resonance (SPR) experiments were conducted using a Biacore T200 instrument with Series S CM5 sensor chip (Cat. BR10053014100530, Cytiva). All experiments were conducted in HDS-EP buffer (Cat. BR100826, Cytiva) at 25 °C. The IW9-pHLA complex served as the analyte, while the TCRs were immobilized as ligands. For steady-state experiments, each TCR was immobilized on the CM5 sensor chip at a density of approximately 1000 response units. The IW9-pHLA complexes were then flowed over all four flow cells at a rate of 10 μL/min. The binding interaction between TCR and pHLA was measured across a concentration range of 0 to 200 μM IW9-pHLA. Each injection had a contact time of 200 seconds, followed by a dissociation time of 1000 seconds. Data were processed using BiaEvaluation 4.1 software and fitted to a 1:1 binding model. Kinetic parameters were calculated, with the dissociation rate globally fitted to ensure accurate determination of binding kinetics.

### Generation of monoclonal TCRs

Monoclonal TCRs specific for the HLA-B*58:01*-*restricted IW9 epitope were expressed using previously described methods^101,102^. TCRα:β libraries from IW9-specific CD8^+^ T cells, with either high or low avidity ratios, were cloned into a modified pLVX-EF1α-IRES-mCherry vector (Takara Bio, Mountain View, CA). This vector was used to generate lentiviruses for transducing the IW9-specific monoclonal TCRs.

### Lentivirus production

Lentiviruses were produced using the adherent 293FT cell line (Thermo Fisher Scientific, Waltham, MA). Cells were transfected with a plasmid mixture containing the TCR, PSPAX and PMD2G plasmids (Addgene, Watertown, MA), using Lipofectamine 3000 Transfection Reagent (Thermo Fisher Scientific, Waltham, MA). A master mix was prepared by combining the plasmid DNA in a 4:1:2 ratio with 40 µL of P3000 Reagent and 12 µg of the pLVX-EF1a-IRES-mCherry plasmid in 1 mL of Opti-MEM Reduced Serum Medium per T75 flask. Separately, Lipofectamine 3000 was diluted 1:25 in 1 mL of Opti-MEM. The diluted transfection reagent was then combined with the master mix and incubated at room temperature for 15-20 minutes. This mixture was added dropwise to the 293FT cells and incubated for 3 days at 37 °C.

### Virus concentration and transduction

The collected supernatant containing lentivirus was centrifuged at 800 x g for 10 minutes and concentrated using a 3:1 ratio of Lenti-X concentrator (Takara Bio, Mountain View, CA). The concentrated lentivirus, along with 4 μg/mL of polybrene (Sigma-Aldrich, St Louis, MO), was then used to transduce a TCR-null Jurkat cell line (BPS Bioscience Cat. No: 78539). The transduced mixture was incubated overnight at 37 °C with 5% CO_2_.

### Post-transduction processing

Single-pair TCRαβ sequences were cloned into a TCR-null Jurkat cell line to evaluate their antigen sensitivity. This was assessed through the expression of CD69 in a T cell activation assay at varying concentrations of HIV-1 antigens. Following transduction, the cells were centrifuged at 500 x g for 5 minutes, resuspended in 10 mL of prewarmed RPMI 1640 medium supplemented with FBS, L-Glutamine, penicillin and HEPES Buffer. The cells were transferred to T25 culture flasks and incubated for 3 days at 37 °C with 5% CO_2_. After incubation, the cells were washed twice with PBS and sorted by FCAS to isolate mCherry-expressing monoclonal T cells. The sorted mCherry-positive cells were cultured in R10 media for 7 days, after which CD3^+^ cells were isolated using the REAlease CD3 Microbead Kit, (Miltenyl Biotec). Tetramer staining was then performed on the CD3^+^ enriched monoclonal cells.

### Antigen sensitivity assay

Dose response experiments were conducted using *HLA-B*58:01* monoallelic TAP-deficient cells as antigen-presenting cells (APCs). Equal numbers of APCs were pulsed with varying concentrations of peptide and incubated for 4 hours at 26 °C. These peptide-pulsed APCs, referred to as target cells, were then cocultured with the same number of monoclonal T cells in 96-well plates overnight. APCs pulsed with irrelevant peptides were included as a negative control, along with a no-peptide condition. Cell activation was assessed via flow cytometry by staining for CD69 (Cat. 310904, BioLegend) following 18 hours of incubation. Peptide dose-response curves for each TCR were generated to estimate antigen sensitivity, with the EC_50_ serving as a quantitative proxy for TCR functional responsiveness. EC_50_ values were estimated using a 4-parameter logistic model in GraphPad Prism (version 8.0) and reported as pEC_50_ (-log_10_ EC_50_) to allow comparison across TCRs of different avidities.

### Quantification and statistical analysis

Dot plots, nonparametric statistical analyses, multiple comparison corrections, and nonparametric correlations (Spearman’s rank correlation) were conducted using GraphPad Prism version 8.0. Group differences were evaluated using the nonparametric Mann-Whitney U test.

## Supplementary Materials

**Figure S1:**
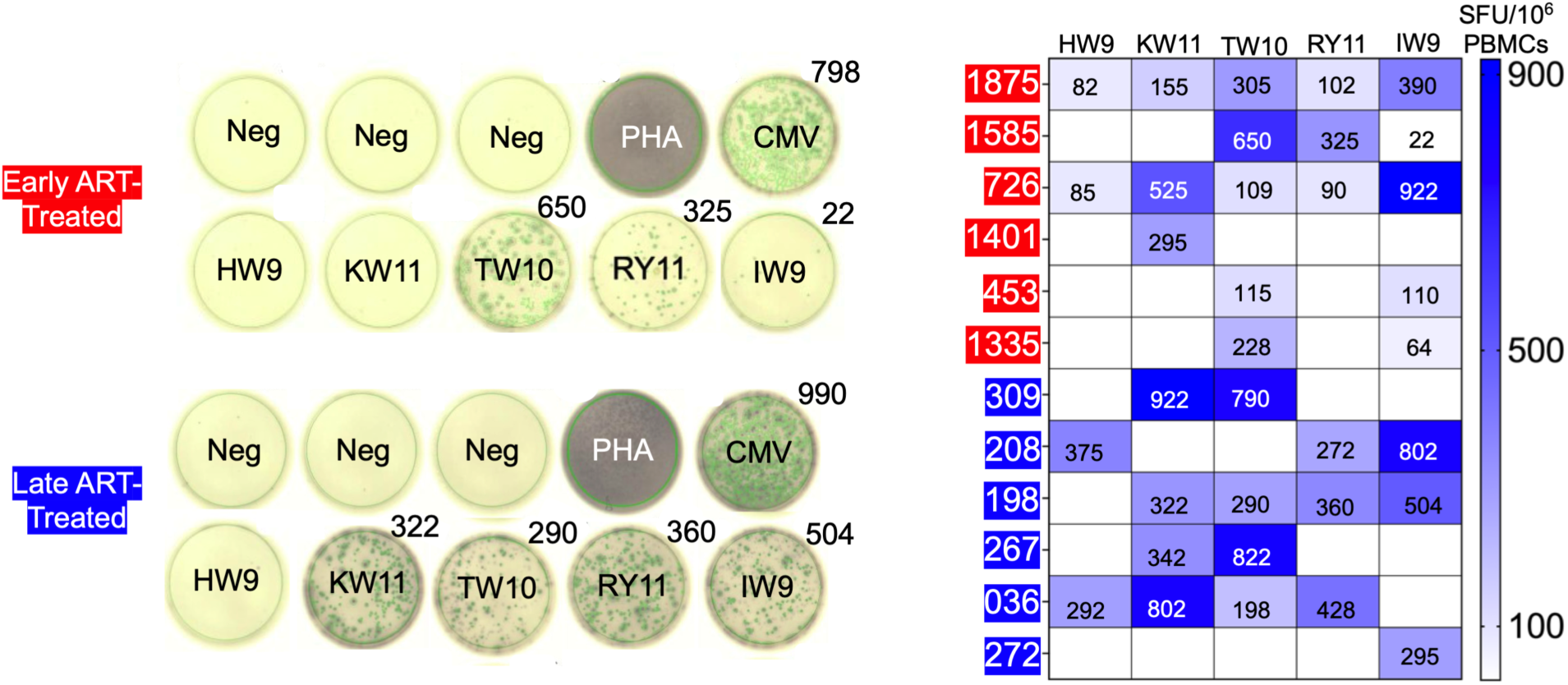
HLA-B*58:01-restricted HIV-specific CTL responses *ex vivo* and following *in vitro* expansion: Breadth and magnitude of HIV-specific CTL responses in PBMCs, measured by IFN-g ELISPOT to five clade C HLA-B*58:01-restricted consensus epitopes (TW10, IW9, KW11, RY11, and HW9) in the 12 participants, measured during the acute phase of HIV-1 infection. Reported values represent spot-forming units (SFUs) after subtraction of background activity from triplicate negative control wells without peptide stimulation^33^. Each column corresponds to an individual epitope, with response magnitudes depicted by a color scale. Numbers within each box indicate SFUs per 10^6^ PBMCs. The intensity scale shown on the right denotes the strength of responses above background. Early and late treated persons are indicated by identifiers shown in red and blue boxes, respectively.

**Figure S2:**
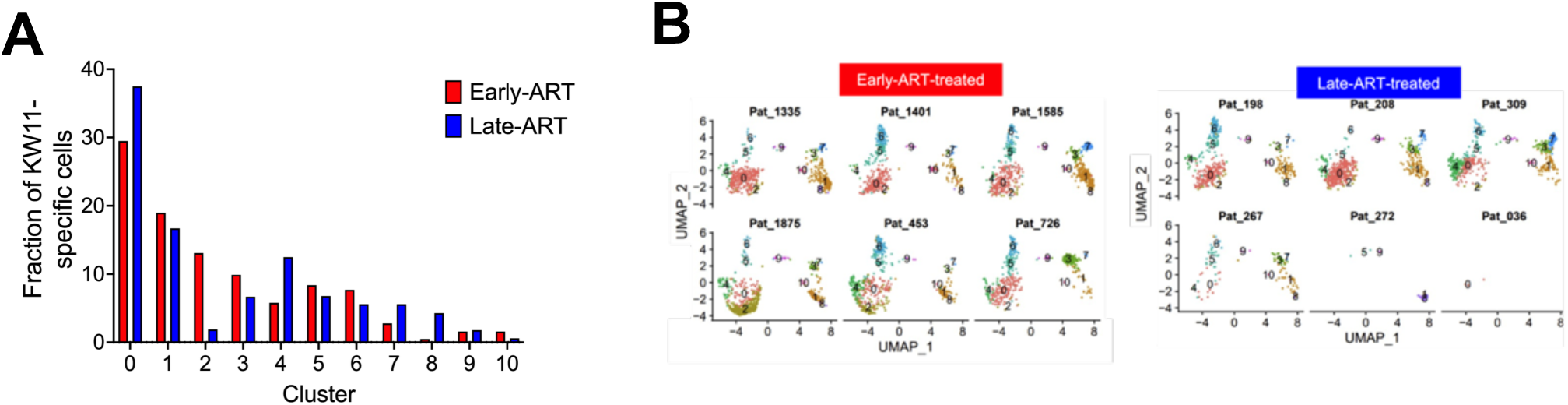
Comparison of KW11-specific clonotypes captured *ex vivo*. (**A**) Bar graphs showing the proportion of KW11-specific CD8^+^ T cells from each cluster in Figure 6B. Red bars represent early ART-treated donors, and blue bars represent late ART-treated donors. (**B**) Cluster distribution by donor, showing the relative fraction of cells per participant, with donors 272 and 036 having limited cell recovery.

**Figure S3:**
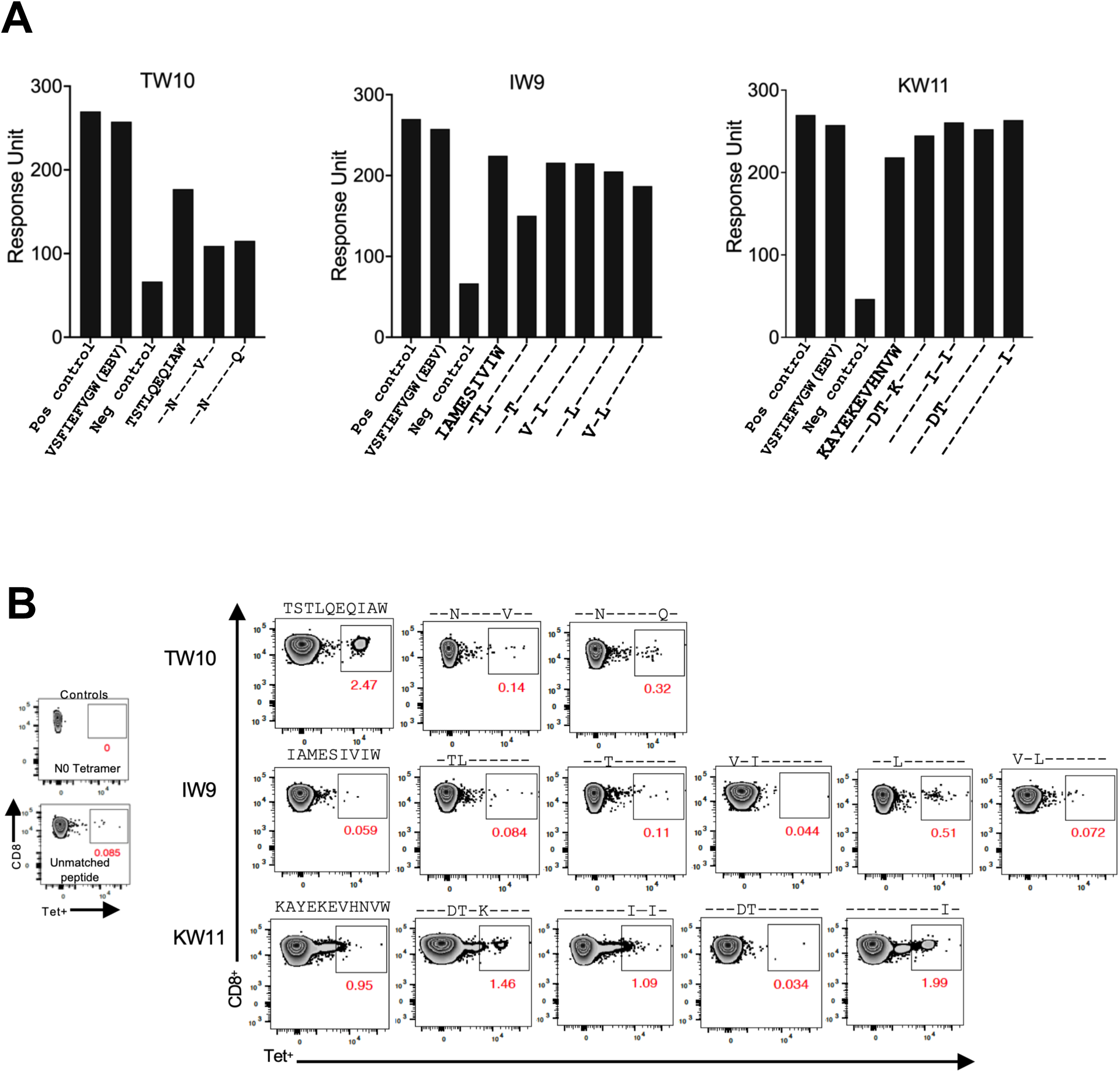
Validation of barcoded tetramer folding using Flex-T ELISA assay: **(A)** The bar graphs show response units from Flex-T sandwich ELISA assay, utilized to verify the correct folding of each barcoded tetramer. This assay confirmed proper folding for most barcoded tetramers, except for the TW10 variants, consistent with the stability results reported in this study. **(B)** Barcoded tetramers were used to stain CD8^+^ T cells following expansion with soluble CD3/CD28, followed by a 2-day resting period. Representative plots are shown from one of the late-treated donors.

**Figure S4:**
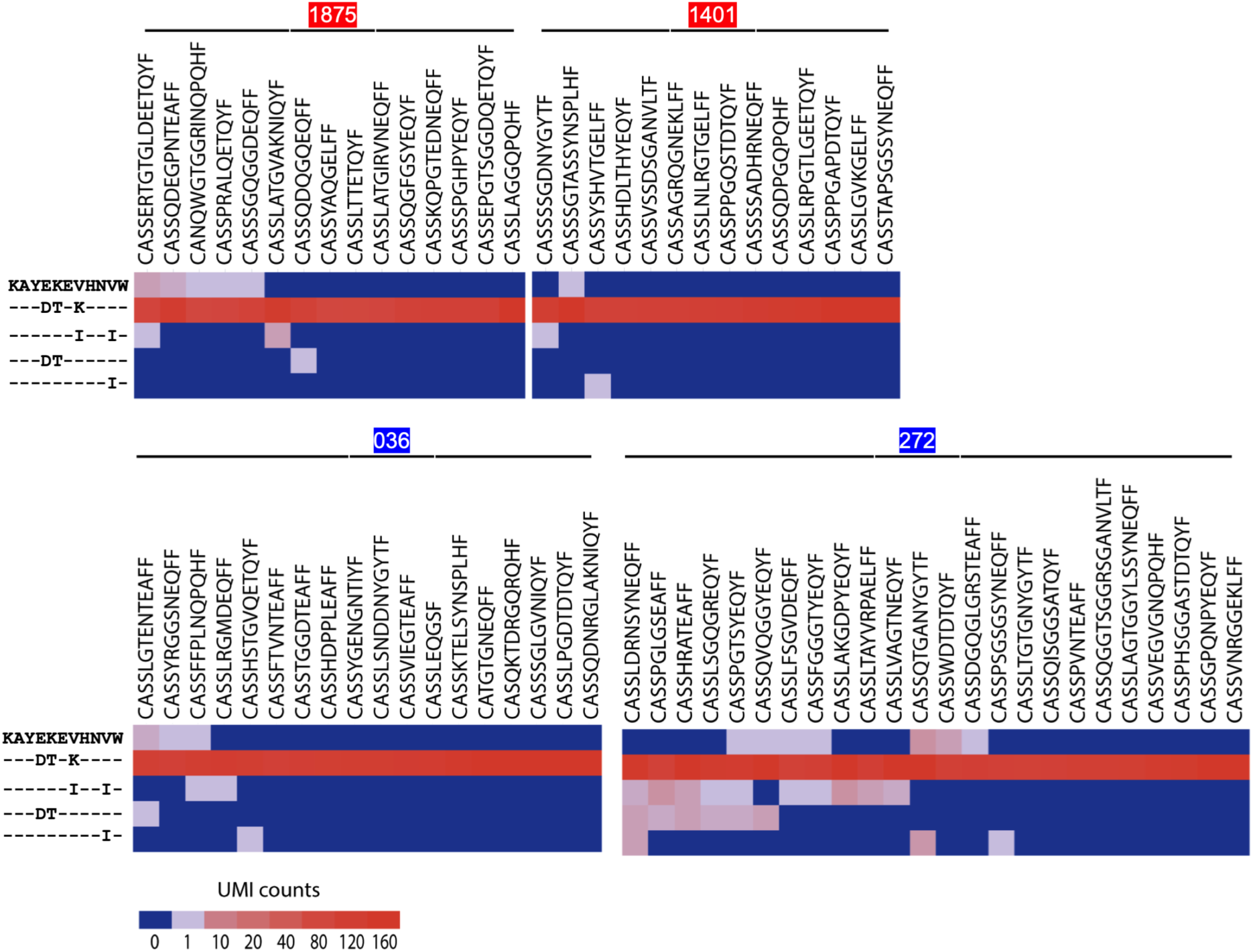
Representative heatmaps from two early treated (1875 and 1401) and two late-treated (036 and 272) participants, illustrating examples of the intensity of tetramer UMI counts per cell, stratified by assigned epitope-specificity. Several of the clonotypes shown here correspond to thise included in the Figure 4. Data are shown for the KW11 epitope using pooled, barcoded pHLA tetramers encompassing both consensus and variant KW11 epitopes. All epitopes used for pHLA tetramers were obtained from HLA-B*58:01-positive donors. Color intensity reflects increasing tetramer UMI counts for each clonotype, annotated by its CDR3β sequence. Clonotypes were considered positive if detected with ≥5 counts.

**Figure S5:**
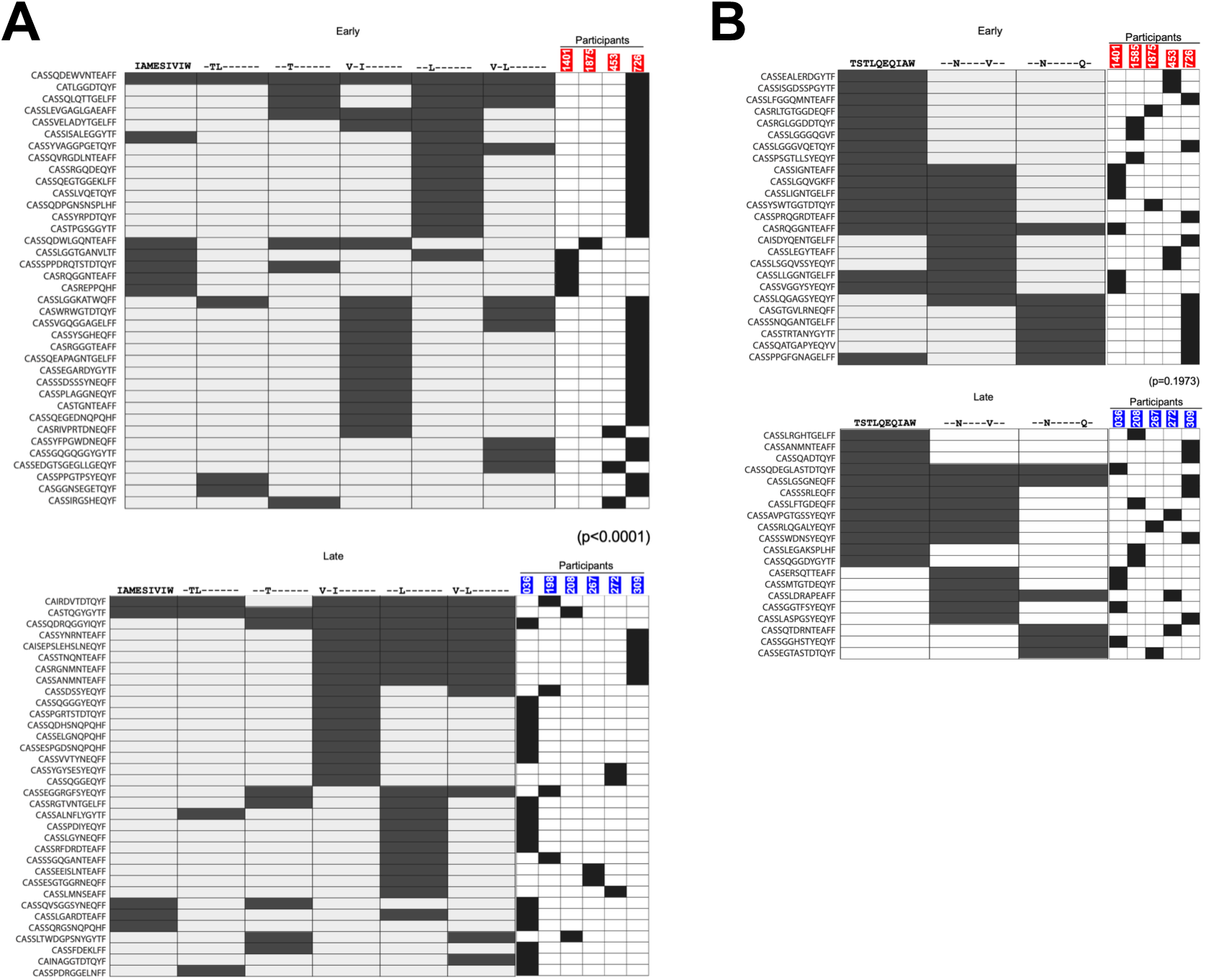
HIV-specific TCRs recognition in early versus late treated groups. Heatmaps illustrating cross-reactive TCR clonotypes targeting (**A**) the IW9 and (**B**) the TW10 antigen family. Clonotypes were detected in donors with immediate (top) and delayed ART-treatment initiation. Each heatmap shows TCR recognition and binding strength across consensus and variant epitopes. The right column indicates the relative contribution of individual donors within each treatment group to the observed cross-reactive pattern. For the IW9 epitope, 66% of IW9-specific clonotypes from early ART-treated individuals were monospecific, compared with 55% in late ART-treated individuals. For the TW10 epitope, 57% of TW10-specific clonotypes from early ART-treated individuals were monospecific, compared with 60% in late ART-treated individuals are not shown in these monochromatic panels.

**Figure S6:**
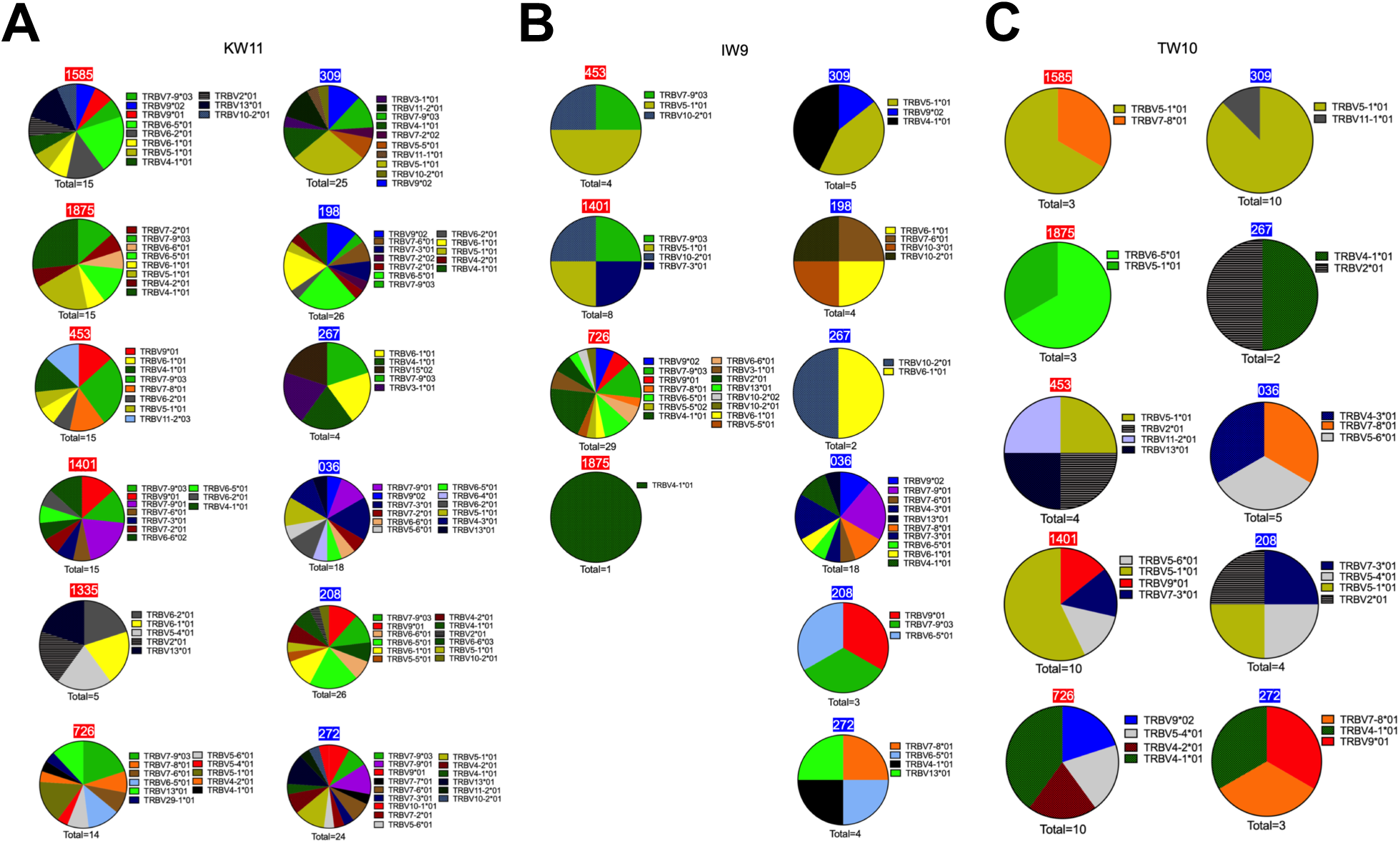
**Shared TRBV gene usage among HIV-specific TCRs from early and late ART-treated individuals**. (**A**) KW11, (**B**) IW9, and (**C**) TW10-specific clonotypes are shown. Each pie chart summarizes the proportion of TRBV genes usage within individual donors, with columns grouped by treatment category for each epitope. Donor IDs are indicated above each pie chart, with early ART-treated donors shaded in red and late ART-treated shaded in blue. The “total” value below each pie chart denotes the number of unique clonotypes detected. Pie slice size represents the relative frequency of clonotypes using each TRBV gene within that donor, with color coding to the right for each pie chart.

**Figure S7:**
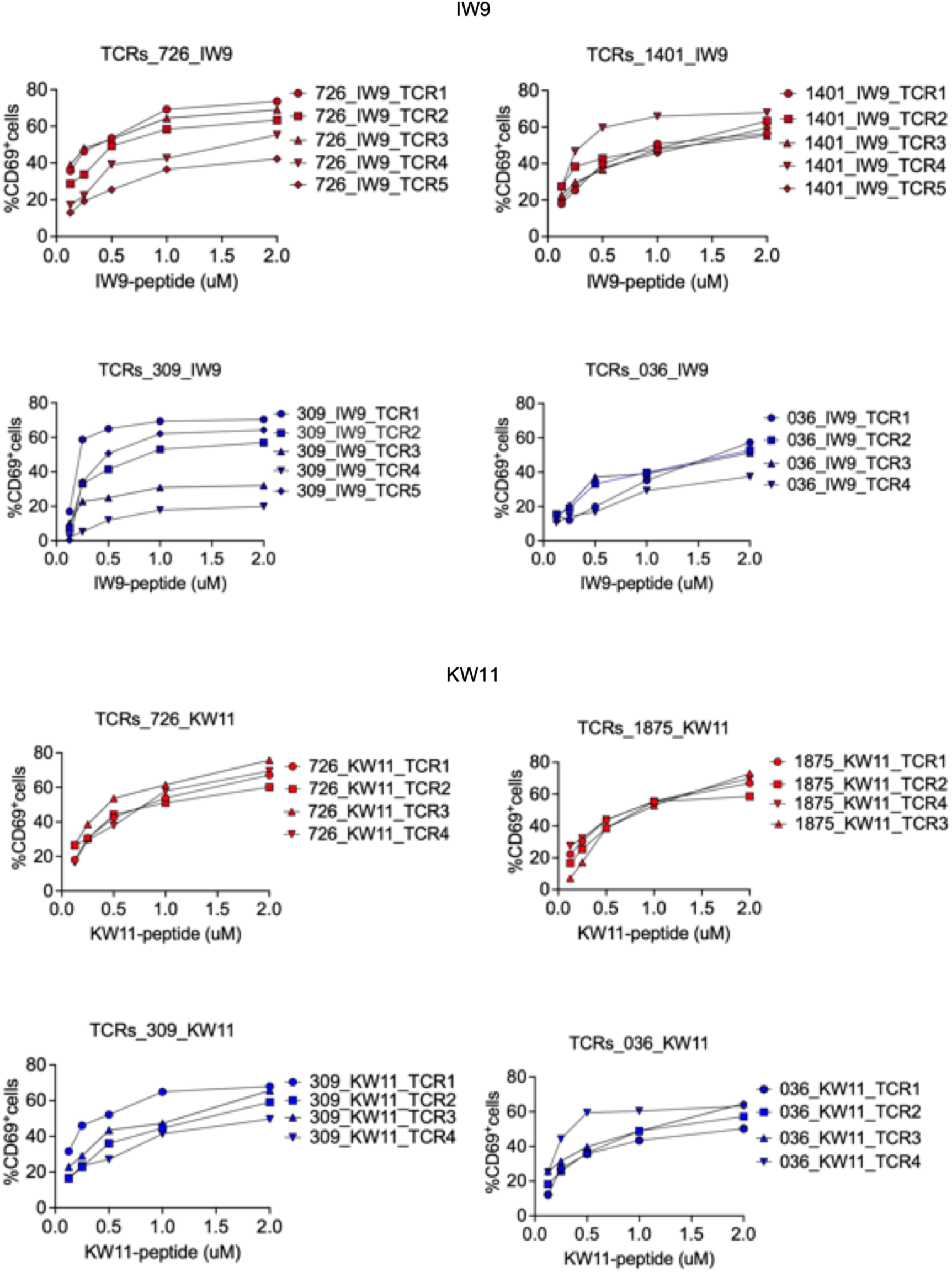
Normalized dose-response curves showing CD69 upregulation in monoclonal TCR-expressing Jurkat cells stimulated with IW9 (top) or KW11 (bottom) peptides. The Y-axis indicates the % of CD69^+^ Jurkat cells, and the X-axis represents peptide concentration. Each curve corresponds to an individual TCR, labeled by its respective ID. Red curves denote TCRs derived from early-treated donors, while blue curves represent TCRs from late-ART-treated donors. Peptide dose-response curves for each TCR were generated to estimate antigen sensitivity, with EC_50_ serving as a quantitative proxy for TCR functional responsiveness.

**Figure S8:**
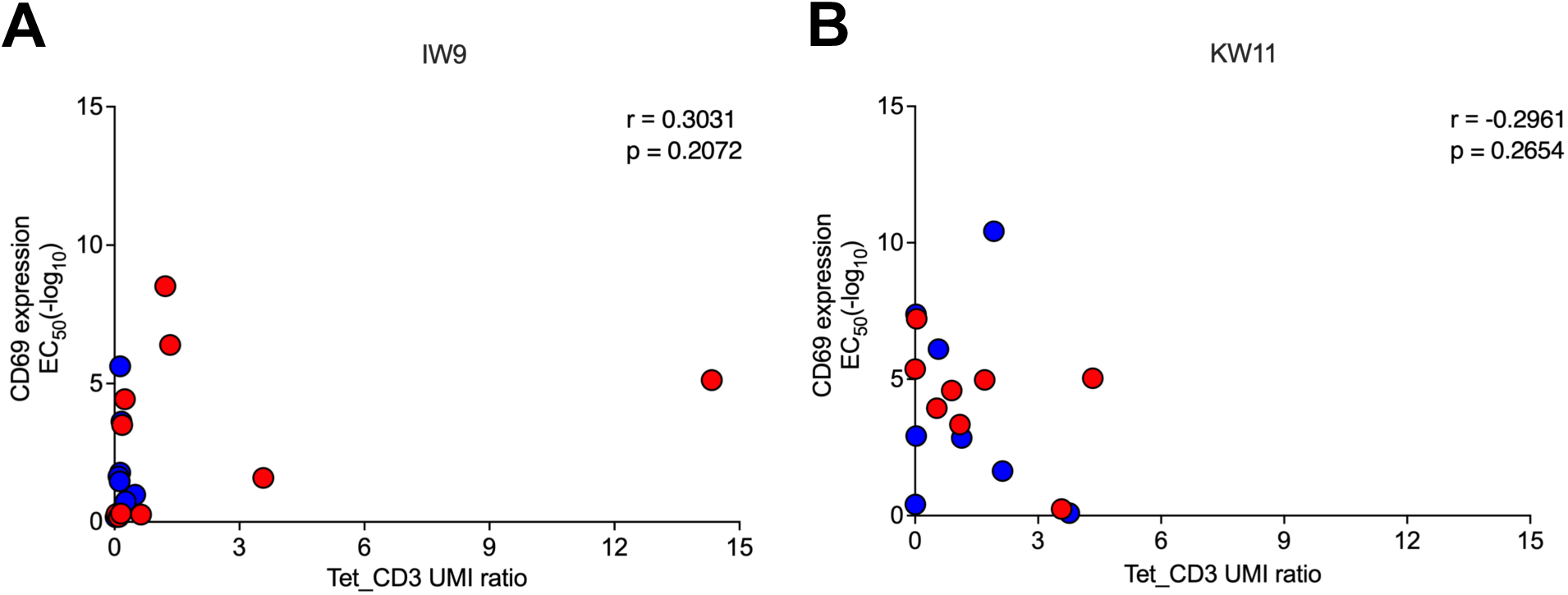
Relationship between peptide dose-responses and tetramer/CD3 UMI ratio. (**A**) IW9 and (**B**) KW11-specific TCRs. The EC_50_ for CD69 expression profile from peptide dose-response assays for each clonotype is on the y axis, and the corresponding tetramer/CD3 UMI ratio on the x-axis. Red dots indicate early ART-treated donors, and blue dots indicate late ART-treated donors.

**Supplementary Table 1:** HLA class I genotype of study participants.

| Participant ID | HLA class I type |
| --- | --- |
| 1401 | A*2,29; B*15,58:01; Cw*3, |
| 1335 | A*2,74; B*50,58:01; Cw*6,7 |
| 1585 | A*23,30; B*15,58:01; Cw*6,18 |
| 453 | A*1,30; B*42,58:01; Cw*6,17 |
| 1875 | A*2; B*45,58:01; Cw*7,16 |
| 726 | A*30,33; B*53,58:01; Cw*4,6 |
| 267 | A*23,74; B*35,58:01; Cw*4,6 |
| 309 | A*2; B*58:01; Cw*7 |
| 208 | A*23,68; B*8,58:01; Cw*3,7 |
| 198 | A*23,30; B*15,58:01; Cw*3,16 |
| 036 | A*2,30; B*44,58:01; Cw*4,6 |
| 272 | A*30,68; B*18,58:01; Cw*2,3 |

## Acknowledgements

We thank the FRESH study participants for their generous contributions. We also thank the FRESH and HIV Pathogenesis Programme staff for their invaluable support in sample collection and processing. Special thanks to Dr. Shiv Pillai for his insightful feedback during the preparation of this manuscript.

## Funding

This study was supported by funding from the Howard Hughes Medical Institute, the Harvard Center for AIDS Research (CFAR) grant (P30 AI060354), the Mark and Lisa Schwartz Foundation, the Philip and Susan Ragon Foundation, the Witten Family Foundation, and the Sub-Saharan African Network for TB/HIV Research Excellence (SANTHE), funded by the Science for Africa Foundation (Del-22-007) with support from Wellcome Trust and the UK Foreign, Commonwealth & Development Office, as part of the EDCPT2 programme supported by the European Union. Additional support was provided by the Bill & Melinda Gates Foundation (INV-033558); and Gilead Sciences Inc. (19275). This work was also supported in part by an NIH Director’s New Innovator Award (DP2-AI158126) to M.E.B., and by the Koch Institute Support (core) Grant P30-CA14051 from the National Cancer Institute. F.J.O. received additional support from the Harvard CFAR. A.R.M received support from the Koch Institute and the Mazumdar-Shaw International Oncology Fellowship. The content contained within is that of the authors and does not necessarily reflect positions or policies of any funders.

## Author Contributions

F.J.O. and B.D.W. conceived the study. F.J.O., N.K.S., B.J.D., M.E.B. and B.D.W. designed the experiments. F.J.O. and N.K.S. performed the majority of the experiments. F.J.O., N.K.S. Z.H. and L.V. performed Immunological assays and 10X experiments. V.B. carried out bioinformatics analysis, while F.J.O. and N.K.S. performed additional statistical analysis. K.R., K.G., and O.O.B. performed viral sequencing. F.J.O., A.R.M., Z.H. and A.F. performed lentiviral generation and activation assays. F.J.O. and N.K.S. performed surface plasmon resonance assays. F.J.O., N.K.S. and A.N. performed peptide stability and thermomelt assays. F.J.O. and Z.H. conducted tetramer binding assays. B.D.W. and M.E.B. supervised the study. F.J.O. and B.D.W. wrote the manuscript with input from T.N. and M.E.B. All authors reviewed the final paper.

## Competing Interests

M.E.B. is a founder, consultant, and equity holder of Kelonia Therapeutics and Abata Therapeutics and received research funding from Pfizer Inc. unrelated to this work. The other authors declare no competing interests.

